# Logistic regression for estimating functional effects with spatial transcriptomics

**DOI:** 10.1101/2025.06.11.659209

**Authors:** Michael Barkasi, Cody Nhan Pham, Demetrios Neophytou, Hysell V. Oviedo

**Affiliations:** Washington University; CUNY Graduate Center

## Abstract

Spatial transcriptomics (ST) unlocks new potential for studying gene functions in cellular processes, as these functions often depend on the orchestration of transcription across space. However, despite their growing number, analysis tools for ST remain largely aimed at data exploration, with few resources for theory-driven hypothesis testing. What’s missing is a way to test whether a factor of interest affects functionally relevant parameters of a gene’s spatial distribution. We present a tool to fill this gap, which we call a warped sigmoidal Poisson-process mixed-effects (WSP, pronounced “wisp”) model. WSP models are the first ST tool allowing researchers to test biologically critical questions without bespoke preprocessing pipelines for identifying key spatial parameters. By aligning coordinates to an axis of interest and letting a likelihood-based regression find between-group effects on expression rates and boundaries, WSP models replace error-prone manual preprocessing with minimally biased hypothesis testing. Integration with genome databases such as GO and KEGG is straightforward, as WSP model estimates of effects take the form of log fold change values. Using MERFISH data collected from wild-type mouse pups, we demonstrate utility by using a WSP model to test the hypothesis that there are interacting effects of age and laterality on gene expression during whisker barrel development.

## 1 Introduction

The spatial distribution of gene expression is a factor in successful biological and cognitive functioning [1–5]. Identifying factors affecting functionally relevant spatial distributions is thus important for understanding the mechanisms behind such success. However, there are few statistical methods suitable for this purpose. Here we address this need by presenting a novel computational tool for estimating effects on the spatial distribution of gene expression. To demonstrate its utility, we use it to analyze effects of age and laterality on the mouse whisker barrel system.

The mystacial vibrissae are the mouse’s largest and most intricately organized tactile organs, providing rich sensory coding central to how the animal explores and engages with its environment [6–8]. A cardinal feature of the whisker-processing cortex is the barrel field in primary somatosensory cortex (S1), where each vibrissa is represented by a discrete, topographically ordered “barrel”. This map forms early after birth and remains highly plastic, reshaping itself in response to sensory experience [9].

The development and maintenance of this topographic organization is orchestrated by a localization of the transcription factor RORβ in cortical layer 4 (L4) of S1 [10, 11]. Given the importance of this localization, it’s important to know which factors *ξ* — such as age, sex, environmental stimuli, or the expression of other genes — have an effect *β*_*ξ*_ on not only the level of RORβ expression, but also on the spatial distribution of that expression across the cortex. We call such effects *functional spatial effects* (FSEs, figure 1A).

**Figure 1.**
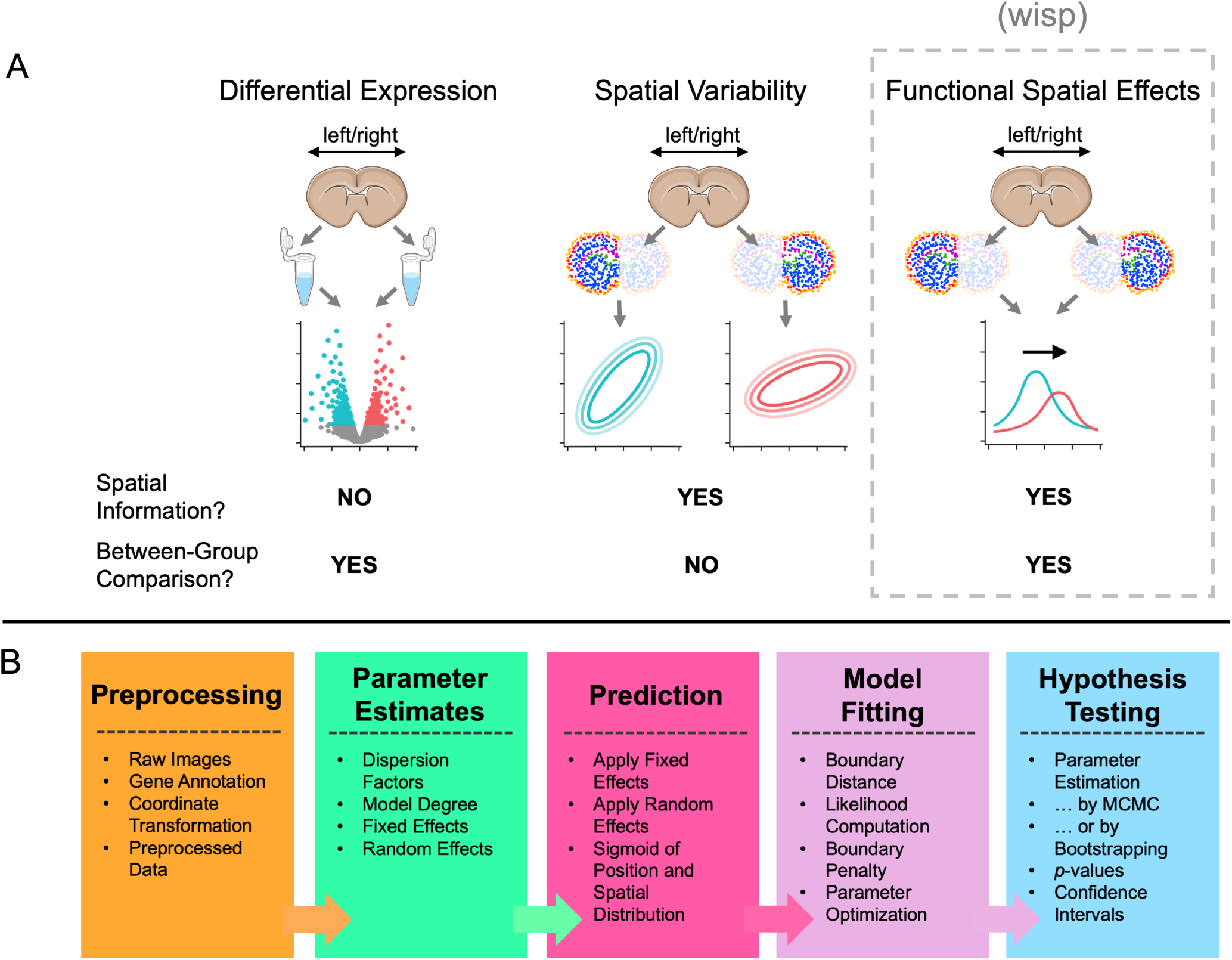
(A) Schematic difference between analyses of differential expression, spatial variability, and functional spatial effects. A depiction of laterality is used as an example to represent a between-group factor. Shown from left-to-right are schematics of a volcano plot, spatial covariance plots, and a spatial-density plot. (B) Pipeline for a WSP model, from data preprocessing to hypothesis testing.

The data necessary for testing hypotheses related to FSEs can be collected through spatial transcriptomics (ST) [12, 13]. These methods measure gene expression while preserving spatial information, either by detecting transcripts directly *in situ* via hybridization (e.g., seqFISH [14], MERFISH [15]) or sequencing (e.g., FISSEQ [16], STARmap [17]), or via arrays or microdissection while using *ex situ* next-generation sequencing (e.g., Visium [18], Slide-seq [19]). In addition, new tools for making targeted interventions on the spatial distribution of transcripts are in development as well [20]. However, current computational methods available for analyzing transcript data were not designed to test for FSEs and are limited in their ability to do so.

Many methods for transcript analysis (e.g., DESeq2 [21] and limma [22]) were developed for use with batch and singlecell RNA sequencing (RNA-Seq) data. These methods model expected transcript count *λ*, or some other measure of gene expression, as a function *f* of categorical factors *ξ*:

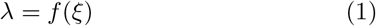

Effects are represented as the difference (i.e., *differential expression*, figure 1A) between a factor’s reference level and treatment level, which for a linear model is the treatment coefficient *β*_*ξ*_. These methods test for effects on *λ* by estimating *β*_*ξ*_. If *λ* is “log-linked”, *β*_*ξ*_ is a log fold change value as used in standard tools such as the gene ontology (GO) database [23] and Kyoto Encyclopedia of Genes and Genomes (KEGG) [24]. New methods for ST data analysis [25, 26], designed to test for spatially variable genes (SVGs, figure 1A), extend the models used in tests for differential expression by including a variable *x* for spatial position:

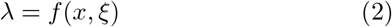

However, testing for FSEs also requires modeling the dependence of the spatial distribution of gene expression on factors *ξ*. If *z* is a variable parameterizing the spatial distribution, such a model would take the schematic form:

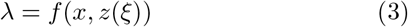

If a coefficient *β*_*zξ*_ is the effect of a factor *ξ* on the parameterization *z* of a spatial distribution of *λ* enabling successful biological or cognitive functioning, then *β*_*zξ*_ would be a FSE. While coefficients *β*_*zξ*_ are not necessarily interpretable as log fold changes, they can be, given the right form for *f* and a log-link.

When the grain of spatial variation is coarse, this modeling can be done by treating the spatial parameterization as just another categorical factor, i.e., *z* = *ξ*, without explicit representation of position *x*. For example, effects on the spatial distribution of RORβ could be estimated by treating L4 as the treatment level of a position factor in a standard model of differential expression with the schematic form of equation 1. However, this approach requires manual preprocessing of the data with a risk of introducing bias; to define the factor *ξ*, the boundaries of the relevant regions of interest (ROIs) must be specified by hand. Further, it’s plausible that many FSEs will involve spatial properties, such as spatial gradients *∂λ/∂x* of expected gene expression, which depend on explicit representation of position and so cannot be represented as categorical factors [27]. In these cases, testing whether the treatment level of a factor *ξ* has a FSE will require a more complex model with the form of equation 3.

Here we present a model of this form, what we call a *warped sigmoidal Poisson-process mixed-effects model* (WSP model, pronounced “wisp”, figure 1B). We have also made available an R package, *wispack* (pronounced “wisp package” or “wisp pack”), which implements WSP models in Rcpp. A minimal working example showing how to use wispack is given below (appendix A.1).

By explicitly parameterizing the spatial distribution of gene expression, WSP models not only allow for testing for FSEs on spatial properties requiring a continuous variable, such as gradients, but also eliminate the need for manual parsing of ROIs. Additionally, WSP models provide estimates of effects on expression rate *λ* in the form of log fold changes. To demonstrate the potential of WSP models, we used one to test for effects of age and laterality on RORβ expression in S1 of wild-type (WT) mice. RORβ is well-suited for this demonstration because it has a known spatial distribution and known functional effects.

We hypothesized that age and laterality would have significant effects on RORβ expression. This hypothesis was based on (1) our previous observations of interacting effects from these factors on electrophysiological recordings of the developing auditory cortex (ACx) of mice [28–30] and (2) histological observations by other labs suggesting the possibility of lateralized development in S1 as well [10, 11, 31]. We tested this hypothesis by fitting a WSP model to transcript counts for RORβ and five other genes known to have layer-specific expression (table 1) collected with MERFISH from male WT mice. As the aim was mainly to demonstrate WSP models in the context of an interesting biological question with real data, we avoided making and testing specific predictions about the size or direction of these effects. However, our concluding discussion outlines interesting ways the FSEs estimated by the WSP model potentially inform current work on RORβ and rodent whisker-barrel development.

**Table 1:**
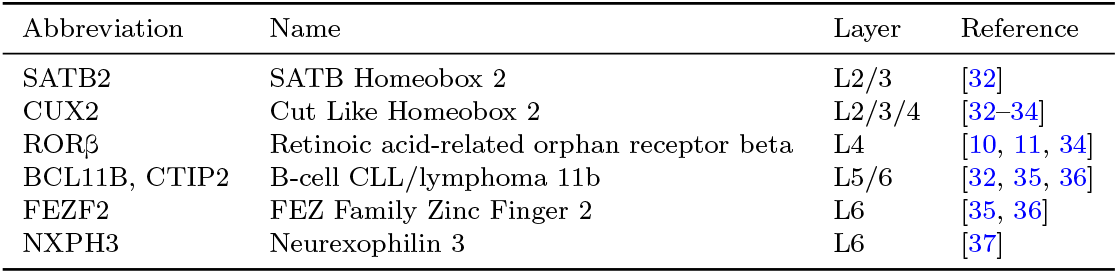
Modeled layer-specific genes and their known layers of expression in the neocortex.

## 2 Modeling methods

### 2.1 Transcript count distribution

Transcript production in a single cell is a stochastic process varying between periods of quiescence and active bursting [38–40]. This variation leads to observed transcript counts *y* with an over-dispersed Poisson distribution. Consistent with normal practice [26, 41, 42], WSP models handle this stochastic behavior by assuming *y* is from a Poisson distribution:

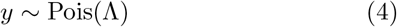

with a stochastic rate Λ from a gamma distribution, i.e., a gamma *kernel* :

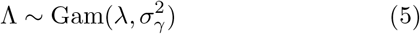

The gamma kernel is parameterized in terms of its expected value *λ* and variance 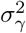, as these are interpretable in terms of expected transcript count, Λ. While Λ is not itself directly observable, it is some unknown random gamma-distributed deviation with variance 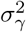 from *λ*. In the standard parameterization of the gamma distribution in terms of “rate” (*γ*_rt_) and “shape” 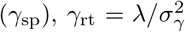 and 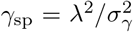. For clarity, we refer to *λ* as the *kernel rate* and Λ as the *transcription rate*. Also for clarity, table 2 provides a glossary of the mathematical notation used here.

**Table 2:**
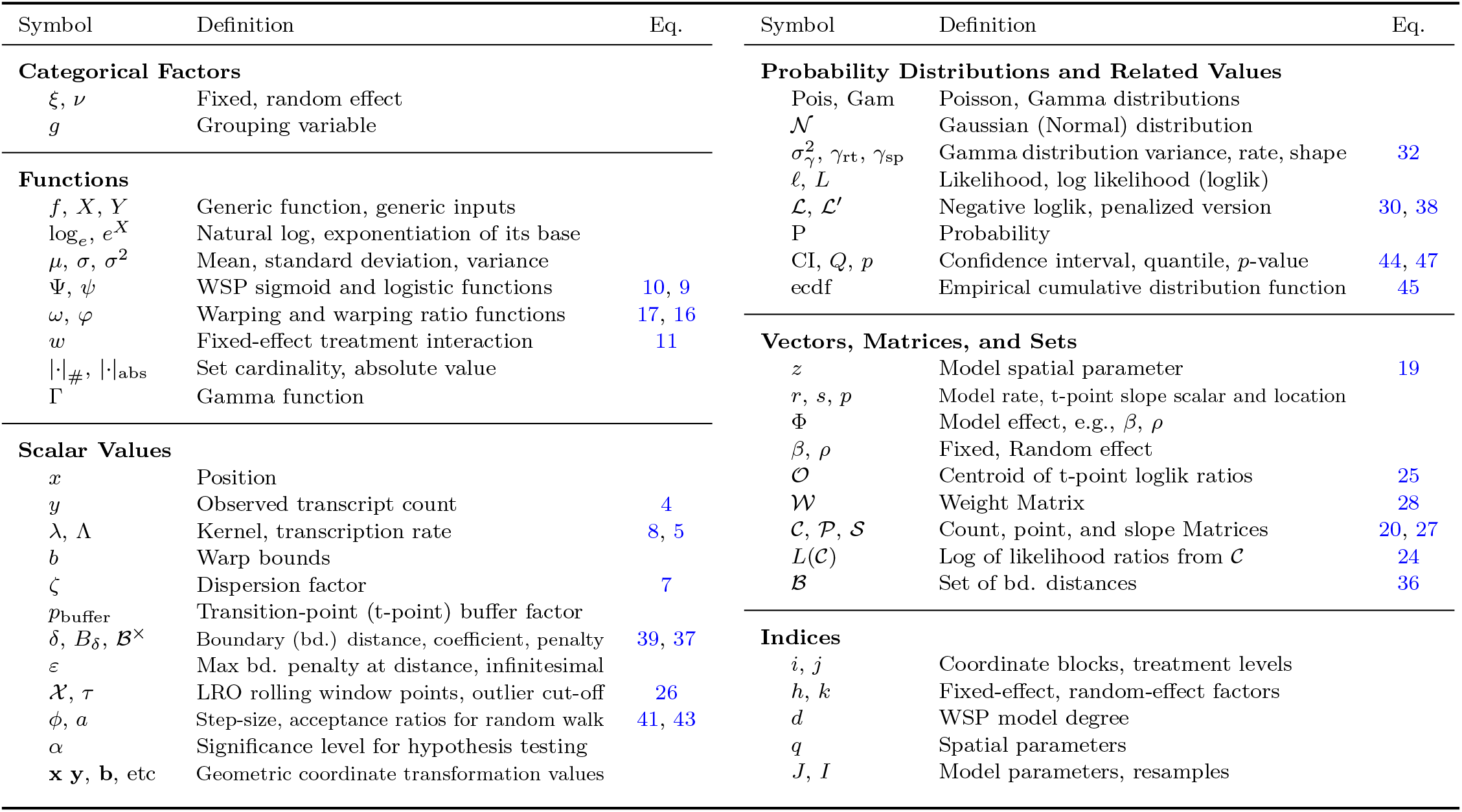
Glossary of notation.

To a first approximation, the aim of fitting a WSP model is to estimate a kernel rate *λ* and variance 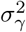 which maximize the likelihood 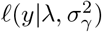 of the observed counts. The kernel rate *λ* is estimated via a prediction from a function *f* of spatial position. Following equation 3, *f* is a function of both spatial position *x* and some parameterization *z* of the distribution of transcripts across *x* which is potentially affected by factors *ξ*. The heart of the WSP modeling approach consists in this spatially parameterized function and will be explained below (equation 10).

Following other standard approaches [21], we assume the variance 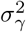 is given in terms of a *dispersion factor ζ* specifying the extent to which *y* is over-dispersed relative to a Poisson distribution:

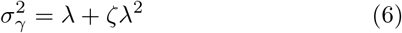

When modeling a set of genes with independent expression rates, the dispersion factor will potentially vary between genes *g*. In this case, *g* becomes a *grouping variable*, a variable the levels (i.e., groups) of which are used to track repeated measures. As our focus is on the prediction of *λ*, we estimate the dispersion factor *ζ*(*g*) with a simple empirical approximation. Specifically, we apply equation 6 to the mean *µ*_*y*_(*g*) (representing *λ*) and variance 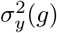 (representing 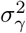) of the observed counts *y* of gene *g*:

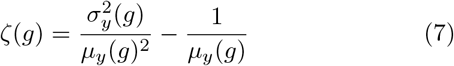

### 2.2 Spatial parameterization

The idea behind WSP models is that gene expression across a spatial region of tissue has a constant kernel rate *λ*, except for transition points where that rate changes. (The transcription rate itself will, of course, vary stochastically from point-to-point.) WSP models capture this idea by predicting log-linked kernel rate for gene expression as a function Ψ of spatial position *x* and three spatial-distribution parameters *z* = ⟨*z*_1_, *z*_2_, *z*_3_⟩, rates *r* = *z*_1_, transition-point slope scalars *s* = *z*_2_, and transition-point positions *p* = *z*_3_:

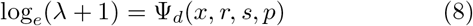

A specific variety of sigmoid, the logistic function (figure 2A), provides the building blocks for the case when *x* is a point in one-dimensional space:

**Figure 2.**
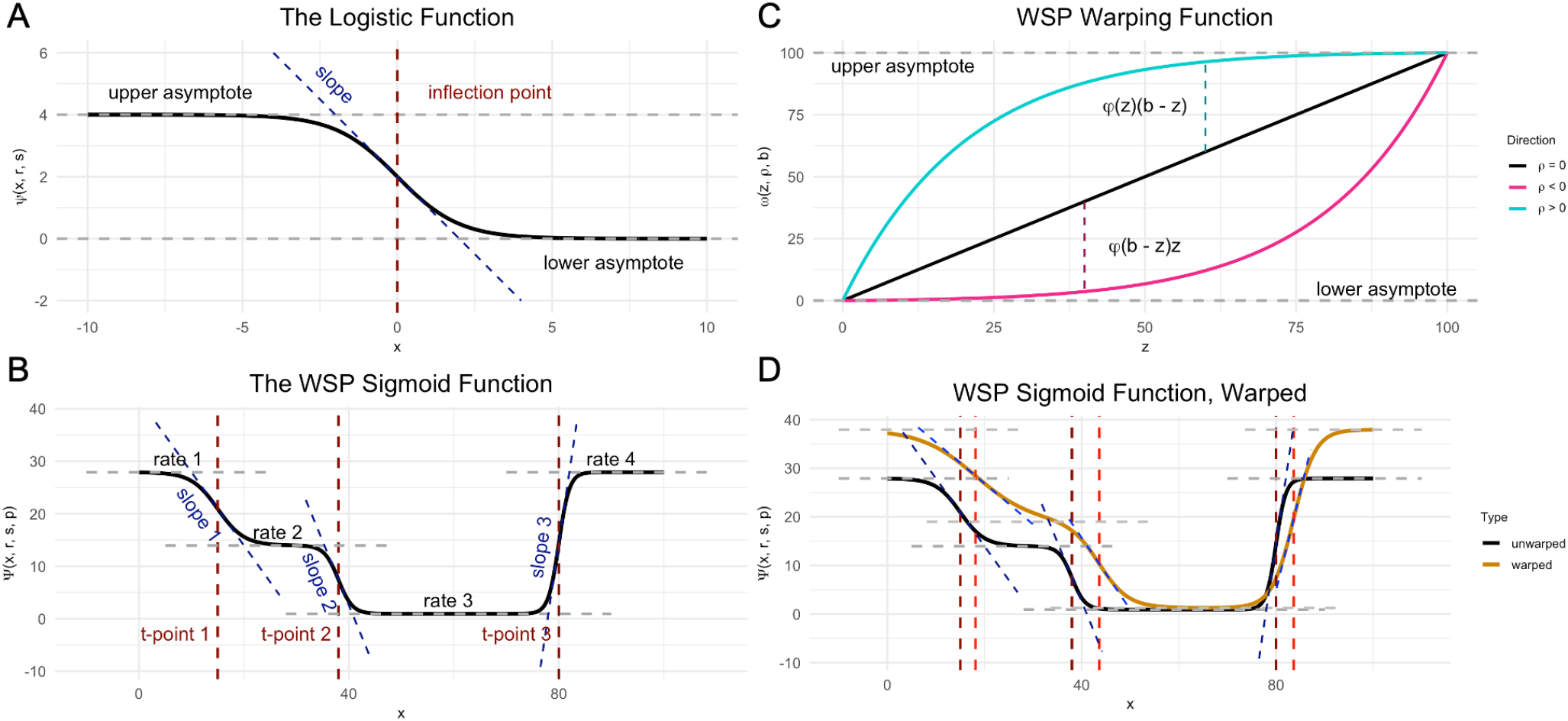
Examples of WSP model and WSP model-related functions, with parameters shown as colored dashed lines. (A) The logistic function. (B) A degree-three WSP sigmoid. (C) The warping function *ω* with positive, negative, and zero warp. (D) A WSP sigmoid warped by the warping function *ω*. Light-colored dashed lines show the displacement of the spatial parameter values under the warp.

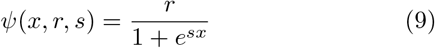

In this equation, *ψ* is a sigmoid with inflection point of slope *s*(*ψr* − *ψ*^2^)*/r* at zero which towards −∞ asymptotes at *r* and towards ∞ asymptotes at zero [43].

While the restriction to one dimension is a limitation, many potential FSEs play out over a single axis, e.g., the laminar axis across the cortex or the radial axis out from a cell nucleus. When using ST data from a tissue slice of negligible thickness, the spatial coordinates can be transformed into a pair ⟨*x, x*^*⊥*^⟩ of orthonormal axes the first of which is defined by the functionally relevant physical axis of interest. If *x* is discretized into bins, the count *y*_*x*_ observed at some position *x* would be the sum of all transcripts within bin *x* with any coordinate *x*^*⊥*^. The additional variance from this aggregation becomes just another source of over-dispersion modeled by the gamma kernel.

Logistic functions are useful for modeling a variable with the features of *λ* because of their behavior when summed, a behavior which, to the best of our knowledge, has not been exploited in regression modeling. Assume *s* and *p* are real-valued vectors of length *d* and assume *r* is a real-valued vector of length *d* + 1 and define, for *d >* 0:

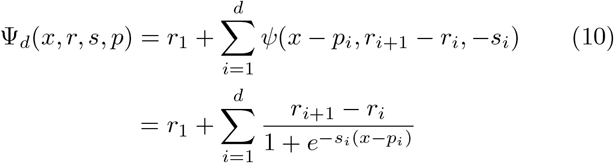

Call *d* the *degree* of Ψ_*d*_ and define Ψ_0_ = *r*_1_. This function maps inputs *x* to constant outputs *r*, except for *d* points *p* at which *r* transitions with slope scaled by *s* (figure 2B). Specifically, using *i* to index the elements of *r, s*, and *p*, if *s*_*i*_ ≫ 0 for all *i* and *i*′ is the largest *i* such that *x* ≫ *p*_*i*_, then Ψ_*d*_(*x, r, s, p*) ≈ *r*_*i*_′_+1_ (appendix A.2). Hence, the value *r*_*i*_ can be thought of as the predicted rate for the block of points *x* such that *p*_*i*−1_ *< x < p*_*i*_, with P_0_ = 0 and P_*d*+1_ = *b*_*p*_. In addition, it can be shown that the slope *∂*Ψ_*d*_*/∂x* at *x* = *p*_*i*_ is approximately (*r*_*i*+1_ − *r*_*i*_)*s*_*i*_*/*4 (appendix A.3). Note that a transition point with a lower slope scalar does not necessarily have a more shallow slope, as the magnitude of the slope can go up even as the scalar goes down, provided the magnitude of the rise *r*_*i*+1_ − *r*_*i*_ increases proportionately more.

All three spatial parameters have a lower bound of zero. Log-linked rate *r* cannot go below zero because of the added one (equation 8), as that would entail negative transcript counts. The slope scalar *s* does not control the sign of transition point slopes (which is controlled by the rise) and, as implied by the sigmoid rate theorem (appendix A.2), cannot go below zero without disrupting the relationship between Ψ and *r*. While the spatial coordinate *x* and the transition points *p* can be any real value, we assume for simplicity that the modeled axis *x* is finite and bound by zero and some upper limit *b*_*p*_.

Each independent gene is modeled with its own function Ψ_*d*(*g*)_, with *d*(*g*) being the number of kernel-rate transition points for *g* — loosely speaking, the number of expression-rate transition points. The estimation of each gene’s degree *d*(*g*) is performed in an initial step before refining the fit of Ψ_*d*(*g*)_ by gradient descent. However, it will be easier to explain this estimation after introducing fixed and random effects.

### 2.3 Fixed effects

The aim of WSP models is to test for *fixed effects* on gene expression, i.e., effects from systematic, reproducible factors *ξ* = ⟨*ξ*_1_, …, *ξ*_*n*_⟩ which divide observations into two classes, a reference class and a “treatment” class. Reference and treatment classes could be defined by a deliberate intervention, e.g., comparing wild-type animals to genetic knock-outs, but could also be any comparison of interest, e.g., comparing old and young animals. These factors are handled just as in a traditional linear model. Each variable *ξ*_*h*_ can take one of two values, zero or one. Hence, *ξ* ∈ {0, 1}^*n*^. The model considers all possible unique treatment interactions 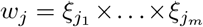, for *m* such that 0 ≤ *m* ≤ *n*. Each treatment defines a weight function *w*_*j*_ : {0, 1}^*n*^ → {0, 1} by multiplication:

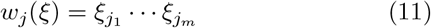

By convention, the first treatment interaction *j* = 1 is the case when *m* = 0, i.e., no treatment. In this case, *w*_*j*_(*ξ*) = 1 for all *ξ* and is the absolute reference level. For each *z*_*q*_ (i.e., for each of *r, s*, and *p*), each such interaction *w*_*j*_ will have, for each group *g*, a corresponding effect value *β*_*qij*_(*g*) for each element *z*_*qi*_(*g, ξ*) of *z*_*q*_(*g, ξ*) such that:

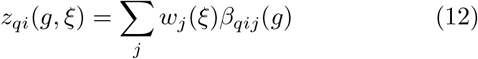

While fully specifying a fixed effect requires specifying *g, ξ, q*, and *i*, we will only write what is necessary for clarity and will also use obvious substitutions, e.g., writing “*β*_*r*_” for *β*_*q*_ when *q* = 1. We adopt this shorthand for not only fixed effects, but all variables.

Given that *r* is, when sufficiently far from the transition points *p*, the log of the predicted kernel rate *λ* plus one (appendix A.2), the rate effects *β*_*r*_ are approximately multiplicative and interpretable as log fold changes. That is, if *β*_*ri*1_, …, *β*_*ril*_ are the rate effects for block *i* for each of the *l* possible treatment conditions *w*_*j*_, then the predicted kernel rate (plus one) of a sample in block *i* is approximately the product:

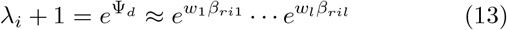

In this equation, *w*_1_ = 1 and the other *w*_*j*_ are zero or one, depending on whether the sample falls under treatment condition *j*. Equation 13 implies that:

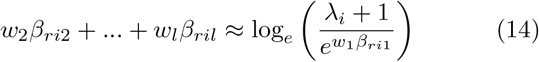

That is, the sum of the rate effects *β*_*r*_, excluding the baseline case of *j* = 1, is the natural-log fold change of the predicted kernel rate *λ* (plus one), the “fold” being with respect to the no-treatment reference condition of *j* = 1. To get the more conventional log-2 fold change value, simply divide this sum by log_*e*_(2). Hence, the log-2 fold change associated with the treatment conditions on kernel rate (plus one) in block *i* is given as:

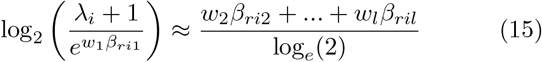

This ability to extract log-2 fold changes makes it easy to incorporate WSP models into a pipeline using standard tools such as Go [23] and KEGG [24].

Transition points *p* and slope scalars *s* are not transformed by the log-link (equation 8), and so the effects *β*_*p*_ and *β*_*s*_ on them are additive. Thus, they cannot be interpreted as log fold changes, but can be read straight off equation 12. Regardless of the log-link, for *j* ≠ 1 effects for rate *r*, transition point *p*, and slope scalar *s* are all interpreted in the usual way: zero is no effect, negative values represent a decrease in value, and positive values represent an increase. Thus, when *j*≠ 1, fixed effects *β*_*qij*_ can be modeled as coming from a normal distribution centered on zero. When *j* = 1, while treated the same as other effects in equation 12, *β*_*qij*_ gives the value of *z*_*q*_ in the reference level and so must follow the distribution of the spatial parameters, i.e., *β*_*qi*1_ *>* 0 and *β*_3*ij*_ = *p* ≤ *b*_*p*_.

### 2.4 Random effects

Hypothesis testing requires distinguishing variation between observations due to fixed effects *β* from variation due to random noise, i.e., *random effects ρ*. Normalization is sometimes used to filter noise, but requires knowing ahead of time which factors *ν* represent true noise and knowing that these factors do not correlate by chance with an effect of interest. For example, analysis pipelines for RNA-Seq data sometimes normalize total transcript count, but this is not recommended when testing for SVGs, as total transcript count can correlate with spatial variance [44]. An alternative approach, *mixedeffects modeling* [45], used in some models of differential gene expression [21] and other areas of genomics [46] and neuroscience [47], is to include the noise variables *ν* (*random-effect factors*) in the model.

WSP models include a single random-effect factor *ν* the levels of which might represent, for example, different biological specimens or different runs of transcript measurement. For this single random-effect factor, WSP models assign a random effect for each of the three spatial parameters, i.e., a random effect *ρ*_*r*_ on rate, *ρ*_*s*_ on transition-point slope scale, and *ρ*_*p*_ on transition-point position. These random effects are relative to a pseudo reference level (“none”, i.e., no random effect) extrapolated from the data by taking the mean of the log-linked count log_*e*_(*y* + 1), observed in each *x* × *g*× *ξ* combination, across all random-effect levels. Random effects are applied to spatial parameters after computing those parameters from the fixed effects (equation 12) but before computing the predicted log-linked count via the WSP model function Ψ (equation 10, figure 1B).

Including random-effect factors in the model allows for estimating the individual contributions of fixed and random effects by modeling the interaction of fixed-effect and randomeffect factors. For each level *k* (taking values zero or one) of a random-effect factor *ν*, a WSP model assigns each level *g* of the grouping variable a random effect *ρ*(*g, k*). The process of fitting the model implicitly estimates the individual contributions of fixed and random effects by, for each *g*, finding the values *β*_*q*_(*g*) and *ρ*_*q*_(*g, k*) which yield the best fit to the observed data.

Traditional linear mixed-effects models treat random effects as just another value to be added in the sum giving the prediction (equation 12). However, for WSP models each *z*_*q*_ has a lower bound of zero and should be smoothly skewed asymptotically towards that bound by negative *ρ*_*q*_. Further, the upper limit *b*_*p*_ of position and transition points imposes a further bound on positive *ρ*_*p*_, such that those effects should also smoothly skew transition points towards this bound asymptotically.

This asymptotic “warping” behavior can be achieved for positive *ρ* by adding to *z*_*q*_ an asymptotically increasing ratio *Φ* of the distance *b* − *z*_*q*_ to the upper boundary, so long as *Φ* scales with *z*_*q*_. Conversely, subtracting from *z*_*q*_ that same ratio *Φ* of *z*_*q*_ itself will achieve the desired warping behavior for negative *ρ*, so long as *Φ* scales with the distance *b*−*z*_*q*_ to the upper boundary (figure 2C). There are many such functions *Φ*, but the following behaves well:

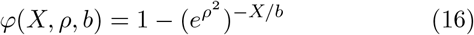

With this equation, the random effects are defined:

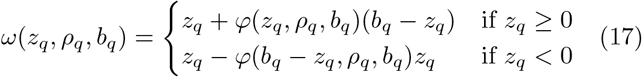

It is easy to see from equation 16 that, when *ρ* = 0, *Φ* = 0 and so *ω*(*z*_*q*_, *ρ*_*q*_, *b*_*q*_) = *z*_*q*_. Thus, as expected, a random effect of zero leads to no warp. Less obviously (see appendix A.4), when *b* → ∞, the warping function reduces to a simple linear function of *z*_*q*_:

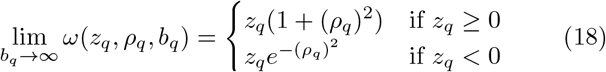

Thus, as expected, when there is no upper bound there is no need to “warp”, i.e., curve, the input. In this case, which applies for rate *r* and slope scalar *s*, random effects *ρ* become linear scaling factors.

Each possible combination *g* × *ξ* × *k* of grouping variable level *g*, fixed-effect factors *ξ* = ⟨*ξ*_1_, …, *ξ*_*n*_⟩, and random-effect level *k* determines, for each spatial parameter *z*_*q*_, a vector of potentially unique values:

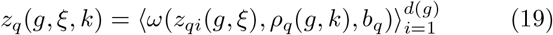

A full WSP model uses this equation to compute the parameterizing inputs *z*_1_ = *r, z*_2_ = *s*, and *z*_3_ = *p* for equation 10 (figure 1B).

### 2.5 Degree estimation

Estimating the degree *d* for each gene *g* amounts to estimating the number of transition points in the kernel rate *λ* along spatial dimension *x*. To this end, we adopt the basic idea behind likelihood-ratio outlier (LRO) change-point detection algorithms [48]. We assume that *d*(*g*) is fixed for each *g*, but that the positions and slopes of transition points potentially vary between both treatment levels 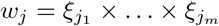 and random-effect levels *k* of *ν*. To estimate a single degree for each gene, the relative likelihood of a transition point compared to the likelihood of no transition point is computed separately for observations across treatment and random levels. These likelihood ratios are aggregated into a centroid and it is this centroid which is tested for transition points by searching for outliers.

Specifically, a matrix 𝒞 is constructed which organizes the observed counts *y* for a gene *g* by spatial coordinates *x* (rows) and the interaction between treatment conditions *w*_*j*_ and random levels *k* (columns). Each column of this matrix 𝒞 gives the total transcript count *y* for *g* in each bin of *x*, observed for a given random level *k* under treatment condition *w*_*j*_. Columns representing a random level not observed under that column’s treatment condition are thrown out.

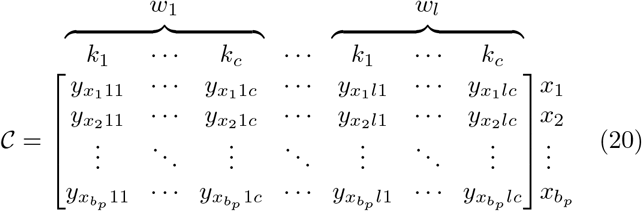

The count values in this matrix are transformed into log space as in equation 8:

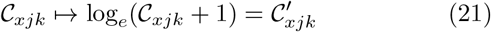

For each column 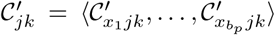 we estimate, for each point 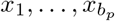, the log of the ratio of the likelihood *ℓ*_cp_ of a change in kernel rate over the likelihood *ℓ*_null_ of no change in kernel rate:

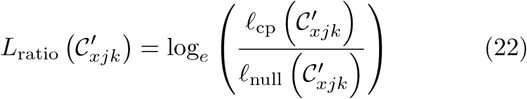

These individual ratios are collected into their own column:

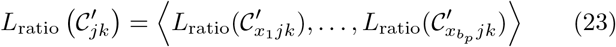

The logs are taken for numerical stability. The likelihood ratios themselves are taken as a measure of how much better a given point *x* is modeled as a transition point vs as a point of constant expression.

The ratios *L*_ratio_ are computed using local windows 𝒳 around the points *x* (appendix A.7). Three different window sizes are used, giving three estimates for each ratio. Multiple window sizes are used because the size of the window corresponds to the frequency of transition points detectable by that window, with smaller windows acting as high-pass filters and larger windows acting as low-pass filters. In wispack, the high-pass window size is a user-defined multiple of the transition-point buffer factor *p*_buffer_ (default in wispack of 1.25) and the mid-pass and low-pass window sizes are 2× and 4× multiples of the high-pass window size.

Next, three matrices *L*(𝒳) of the log-likelihood ratios *L*_ratio_ are defined, one for each filter, as follows:

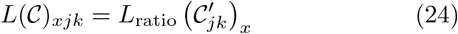

Taking the means across the rows of these new matrices gives, for each of the three filters, an estimate of the centroid 𝒪 of the transition-point log-likelihood ratio over *x* across all combinations of treatment and random-factor levels:

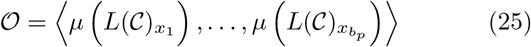

A point *x* is treated as a transition point *p* if and only if, for at least one of the high-pass, mid-pass, or low-pass 𝒪, the element 𝒪_*x*_ in 𝒪 is more than some multiple *τ* of the standard deviation of 𝒪 above the mean:

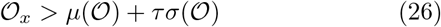

By default in wispack, *τ* = 2.0, on the assumption that transition points rarely make up more than 5% of bins *x*. Potential transition points too close to zero or *b*_*p*_ are thrown out, and if two potential transition points are too close together, the point with the larger of the two log-likelihood ratios is used. “Too close” is defined as within a distance of half the size of the high-pass filter window.

### 2.6 Initial parameter values

WSP models are fit to observations *y* by finding the effects parameters *β* and *ρ* which, given the estimated dispersion factors *ζ* (equation 7), maximize the joint likelihood of those observations within the log-link. This is done by optimizing an initial set of parameter values using gradient descent. Initial values for the random effects *ρ* are picked from a uniform random distribution spanning −0.1 to 0.1. Model fitting is greatly improved if the fixed effects are initialized close to their optimal values, and so the points *p* determined by the LRO outlier threshold (equation 26) are used to set, for each treatment level *w*_*j*_, initial values for all fixed effects *β*_*j*_.

The LRO outlier threshold only estimates a single set of transition points *p* spanning all conditions for a gene, so the first step is to estimate the transition point locations for each combination of treatment and random-effect level. A dynamic time-warping algorithm [49] is used to align each column of the mid-pass *L*(𝒞) to the mid-pass centroid 𝒪 (equation 25). The resulting alignments (one per column) are used to project the points *p* onto the row numbers of *L*(𝒞), yielding one projection 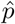 per column of *L*(𝒞). For this step wispack uses the dtw package in R (v1.23-1, [50]) with the warping assuming a symmetric step pattern of slope 0.5 [51]. These projections are collected into a new matrix 𝒞_𝒪_ with the column structure of 𝒞, the number of rows of which is the degree (i.e., is the number of transition points).

The second step is to estimate transition point locations *p* for each treatment level *w*_*j*_ by taking the mean across the rows of 𝒞_𝒪_ within columns under *w*_*j*_. The resulting subsetted and factored matrix has the form:

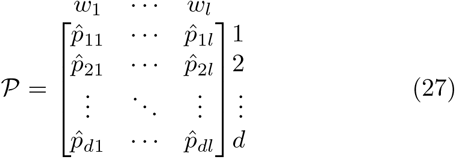

For each transition point index *i* (1 ≤ *i* ≤ *d*), the row 𝒫_*i*_ provides an estimate of the transition point location observed under each treatment condition *j*. These estimates can be used to define *d* + 1 blocks of points *x*, such that block *i* includes 𝒫_(*i*−1)*j*_ ≤ *x* ≤ 𝒫_*ij*_ with 𝒫_0*j*_ = 0 and 𝒫_(*d*+1)*j*_ = *b*_*p*_.

Next, from these blocks a rate matrix ℛ can be defined such that ℛ_*ij*_ is the mean of the log (plus one) of the observed counts *y* in block *i* for treatment *w*_*j*_. Similarly, a slope matrix 𝒮 can be defined such that initial estimates 𝒮_*ij*_ for the slope scalars are calculated by finding the number of bins (points *x*) which must be stepped ahead and behind transition point 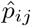 such that the difference of the means of the log (plus one) of the observed counts before and after 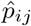 is greater than or equal to some stipulated proportion (by default in wispack, 0.8) of ℛ_(*i*+1)*j*_ −ℛ_*ij*_. This number of bins is the run of the slope, and the implied slope scalar is 4 divided by the run (appendix A.3).

To use the initial estimates ℛ, 𝒮, and 𝒫 of the spatial parameters to set the initial values of the effects *β*_*r*_, *β*_*s*_, and *β*_*p*_, we first define a weight matrix:

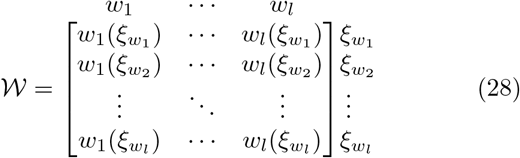

For convenience, 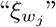 is short-hand for the factor *ξ* which has the components of *w*_*j*_ at treatment level and all other components at reference level. Intuitively, 𝒲 says whether the treatment represented by a column applies (i.e., is weighted 1) in the treatment condition represented by each row. By equation 12, for each transition point or block *i*, treatment level *j*, and spatial parameter matrix *X* (i.e., *X* one of ℛ, 𝒮, and 𝒫), the initial effects *β*_*Xij*_ can be got by solving the following system of equations:

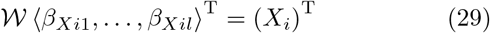

Full-pivot LU decomposition [52] is used in wispack to solve this system.

### 2.7 Parameter optimization

With initial parameters estimated, the next step is to maximize the joint likelihood of the observations *y* within the log-link using gradient descent. It is assumed that (1) genes *g* have independent expression rates, and (2) that once the factors *ξ* and *ν* are fixed, observed count *y*_*x*_ for a gene at position *x* is independent of its observed count at all other positions. For numerical stability, the negative of the log of the joint likelihood is minimized:

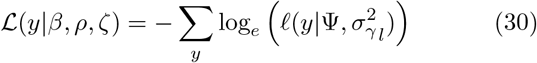

where the likelihood is:

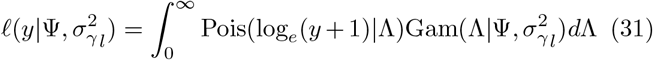

The integral in this equation can be solved analytically (appendix A.5), which greatly simplifies the minimization. Keeping with the interpretation of *ζ* as a dispersion factor (equation 6), if the variable 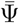 is defined 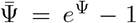, then the variance of the kernel can be approximated using the delta method [53] as:

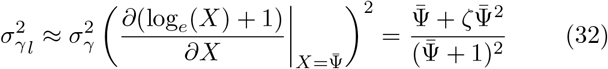

When minimizing the joint log-likelihood of the observations *y* within the log-link, all model parameters must stay within bounds of feasibility. Specifically, (1) all slope scalars *s* must be positive, (2) transition points *p* must be in increasing order, not too close together, and must be between zero and the bound *b*_*p*_, and (3) the predicted value Ψ must always be positive. Regarding this last constraint, Ψ *>* 0 if and only if *r*_*i*_ *>* 0 for all rate components used in the calculation of Ψ (appendix A.6). Thus, it suffices to ensure that all rate components are positive.

To enforce these boundaries, for each possible combination *g*× *ξ*× *k* of *g, ξ*, and *k*, the bounded variables are computed and collected in a set:

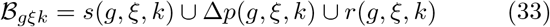

In this equation, *s*(*g, ξ, k*) and *r*(*g, ξ, k*) are the slopes and rates determined by the given combination of grouping variable level, fixed-effects, and random-effect level (equation 19). The middle term Δ*p*(*g, ξ, k*) captures the constraints on transition points. It is defined:

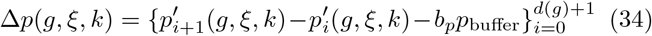

in terms of an extension *p*′ of *p*:

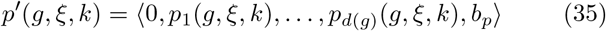

Notice that this equation defines “too close together” as a distance smaller than *b*_*p*_*p*_buffer_, for a scalar *p*_buffer_ between zero and one (set to 0.05 by default in wispack).

The total set of boundary distances is the union of all of these sets:

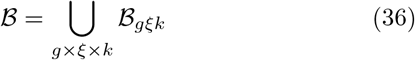

and from this set B a boundary penalty is defined:

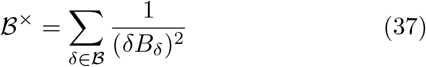

This penalty is nearly zero when all boundary distances *δ* in ℬ are far above zero (*δ* ≥ 1) and smoothly approaches infinity exponentially as any one of those distances approach zero.

It is added to the negative log-likelihood ℒ (equation 30) to obtain the value to be minimized:

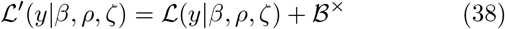

To keep the penalty term B× near zero when all *δ* are far above zero, a weighting coefficient *B*_*δ*_ is included for each distance *δ* such that, for *ε* some near-zero penalty value and the boundary distances computed from the set of initializing effects parameters *β* and *ρ*:

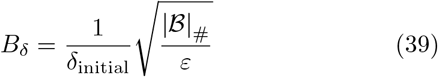

The minimization of ′ is performed in wispack using the LBFGS algorithm (NLopt, [54, 55]) with gradients calculated via reverse-mode automatic differentiation (Stan, [56]).

### 2.8 Hypothesis testing

WSP models test the hypothesis that factor *ξ* has effect *β*_*ξ*_ on gene-expression distribution *z* by estimating the probability of a given value *β*_*zξ*_, conditional on observed transcript counts *y*. That is, the conditional joint distribution P(*β, ρ*| *y, ζ*) is estimated. This can be done by resampling *β* and *ρ*, either (1) using a Markov chain Monte Carlo (MCMC) algorithm, or (2) by refitting the model to resamples of *y* with new estimates of *ζ* (bootstrapping). As fixed *β* and random *ρ* effect parameters are handled the same when resampling, we use the variable Φ to cover both cases and use *J* to index the parameters in Φ.

MCMC resampling is done in wispack with a Metropolis–Hastings algorithm [57] which steps from one parameter vector Φ to the next Φ^+^ by random walk, starting with the same parameters used to initialize the L-BFGS algorithm. Steps are drawn from a normal distribution:

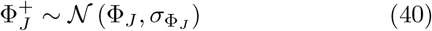

The values in Φ can differ by orders of magnitude and must stay within the boundaries of feasibility. Thus, step size is a fraction *ϕ* of the absolute value of the parameter Φ_*J*_ and further scaled by the boundary penalty 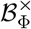 of Φ (equation 37):

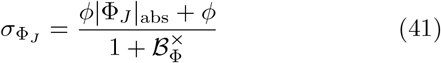

Aside from the reference values *β*_*pi*1_ for the transition points, which are assumed to be uniformly distributed, the prior P_prior_(Φ_*J*_) of each Φ_*J*_ is calculated from the density function of a normal distribution centered on zero with adjustable *σ*. The precise value of *σ* is not crucial, so long as it is around the size of a typical parameter in Φ. By default in wispack, *σ* = 1. The bell shape of this prior captures the assumption that larger values are less likely than smaller ones. The overall log-prior log*e*(P_prior_(Φ)) is computed as the sum of all log_*e*_(P_prior_(Φ_*J*_)). Log-likelihoods *L*(*y*| Φ, *ζ*) are computed according to equation 38 as *L*(*y*| Φ, *ζ*) = −ℒ′(*y* |Φ, *ζ*). The posteriors are computed as:

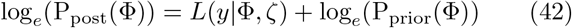

The acceptance ratio *a* for a new step Φ^+^ is then given by:

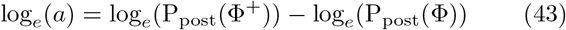

Bootstrapping is viable and preferred if the structure of the data allows for resampling and computational resources allow for refitting. (Bootstraps can be run in parallel by wispack using forking on Mac and Linux systems; wispack does not provide parallelization for Windows.) The data used here provide such an example. In our data, the total count *y*_*x*_ in a bin *x* for gene *g* is the sum of the counts of *g* in each cell in *x*. Thus, the cells themselves can be resampled with replacement and the counts summed again to get a resampling of *y*_*x*_. After resampling every gene *g* in every bin *x*, a WSP model can be fit again to the resampled observations *y*^+^ to get a new parameter estimate Φ^+^. Before estimating Φ^+^ via gradient descent, new disperion factors *ζ* are computed from *y*^+^ using equation 7 and new initial parameter values are computed using the procedure described above, although the original degree estimation is used in all bootstrap resamples.

For both bootstrap resampling and the Metropolis-Hastings random walk, wispack uses a permuted congruential generator (PCG) implemented in C++ for random number generation. PCGs outperform standard algorithms like Mersenne Twister in computational speed, unpredictability, and conformity to expected statistical behavior [58]. The PCG provides sampling from a uniform distribution, which is used directly for bootstraps and transformed into a normal distribution with the Box-Muller algorithm for the random walk.

Whether by MCMC or bootstrapping, resamples *I* yield a matrix 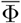 of resampled parameters Φ_*IJ*_. To clarify notation, “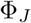” will continue to represent a scalar value for parameter *J*, while “Φ_*J*_ “and” ⟨Φ⟩_*I*_ “represent the column of 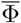 for parameter *J* and row of 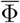 for resample *I*, respectively. Each row Φ_*I*_ is thus a complete parameter vector Φ. A confidence interval for significance level *α* can be computed for each parameter *J* by estimating the first and last (*α/*2)-quantile of column ⟨Φ⟩_*J*_ :

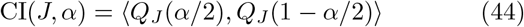

The quantiles *Q*_*J*_ are computed over ⟨Φ⟩_*J*_ with the standard linear interpolation used in R (type 7, [59]).

For a specific observed sample *I*, a *p*-value can be computed from the empirical cumulative distribution function (ecdf), defined for an input vector *X* and scalar value *X*′:

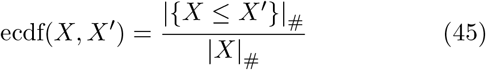

The *p*-value for the observed parameter Φ_*IJ*_, with null hypothesis that Φ_*J*_ = 0, is computed by first recentering the sampled values ⟨Φ⟩_*J*_ around the null:

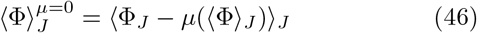

and then estimating the probability of observing parameter *J* with at least the magnitude |Φ_*IJ*_|_abs_, assuming the null distribution:

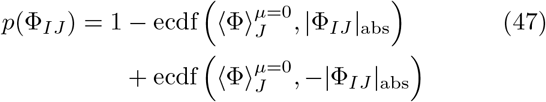

While *p*-values can be computed for any of the resampled parameters, presumably it only makes sense to compute them for some distinguished set of parameters representing the best estimate, e.g., those parameters found by the L-BFGS optimization on the observed data, or an average taken from the resampled parameters. Further, while confidence intervals can be computed for all parameters, the ecdf method (equation 47) only makes sense when the null hypothesis is that the parameter has value zero (presumably representing no effect). Thus, at most the ecdf method makes sense for random-effect parameters *ρ* and fixed-effect parameters *β*_*j*_ such that *j* ≠ 1, i.e., fixed-effect parameters except for the baseline level. The ecdf method works for these parameters because they were constructed to have unconstrained distributions with center zero representing no effect.

In contrast, there is no similarly direct test for a *p*-value from the resamples of the baseline parameters (i.e., *j* ≠ 1), which all have a floor of zero. While there are ways around this difficulty, such as a maximum likelihood ratio test comparing the unconstrained model to one with the target baseline parameter held to zero, these approaches raise new issues, most notably the problem of estimating degrees of freedom. The aim of WSP models is to test for FSEs, not to test whether a given gene is an SVG (non-zero slope scalar) or to test whether that gene is being expressed at all (non-zero baseline rate parameters). Thus, we make no recommendations on how to calculate *p*-values for the baseline parameters. While the ecdf method can be applied to the random effect parameters, wispack does not apply it to them. This is because of the aim of testing for FSEs, but also to limit the number of statistical tests run. Even a handful of genes typically will lead to hundreds of effects parameters in a WSP model, so running additional tests with no clear purpose is not advised. To handle the inevitable multiple comparisons problem with testing even just the fixed effects (for *j* ≠ 1), wispack by default adjusts confidence intervals and *p*-values with a Holm-Bonferroni correction, with the more conservative Bonferroni correction available as well.

## 3 Data collection methods

### 3.1 Tissue collection and MERFISH sample preparation

Male CBA/CaJ mice aged postnatal (P) day 12 and P18 were used in accordance with the National Institute of Health guidelines, as approved by the Washington University in Saint Louis Institutional Animal Care and Use Committee. We had *n* = 2 for each age, for four mice total. Mice were anesthetized with CO_2_ and rapidly decapitated. The brains were harvested, placed in prechilled Optimal Cutting Temperature (Fisher) embedding medium, flash-frozen in 2-methylbutane on dry ice, and stored at −80°C until cryosectioning. Samples were prepared according to Vizgen’s modified formalin fixed-paraffin-embedded (FFPE) protocol for fresh frozen tissue. On the day of cryosectioning, frozen tissue blocks were allowed to equilibrate at −20°C in a cryostat (Leica CM1860 UV) for at least one hour prior to slicing horizontal, 14µm thick sections. Slice were collected on MERSCOPE slides and sections were allowed to adhere to the slides at −20°C for at least 30 minutes before being fixed with warmed 4% PFA in nuclease-free 1X PBS for 60 minutes at 37°C. The samples were then washed in nuclease-free 1X PBS 3× (5 minutes each wash) and air dried in a covered slice rack to ensure complete tissue adhesion. The slices were then incubated in 70% ethanol at 4°C overnight for permeabilization.

### 3.2 Anchoring, gel embedding, and tissue clearing

The following day, sections were washed with conditioning buffer 3× (PN 20300116, 2× at room temperature (RT) for 1 minute, 1× at 37°C for 30 minutes). Sections were then treated with 100µL of Pre-Anchoring Conditioning Buffer (100µL Conditioning Buffer (PN 20300116), 5µL Pre-Anchoring Activator (PN 20300113), 5µL RNase Inhibitor (New England BioLabs)) for 2 hours at RT followed by washes in Sample Prep Wash Buffer (PN 20300001, 1× at RT for 5 minutes) and Formamide Buffer (PN 20300002, 1× at 37°C for 30 minutes). After the formamide wash, the RNA was anchored to the tissue by incubating samples in Anchoring Buffer (PN20300117) for ≈ 16 hours at 37°C. Samples were then washed with Formamide Buffer (1× at 47°C for 15 minutes) and Sample Prep Wash Buffer (1× at RT for 2 minutes), gel embedded (50µL) and cleared (incubated at 47°C for *<* 24 hours). The gel embedding solution contained 5mL of Gel Embedding Premix (PN 20300118), 25µL of 10% ammonium persulfate, and 2.5µL of N,N,N′,N′Tetramethylethylenediamine. Gel coverslips (PN 30200004) were cleaned with RNAseZap and 70% ethanol before 100µL of Gel Slick solution was added. We allowed 15 minutes for the coverslips to air dry thereafter. Clearing solution contained 50µL Proteinase K (New England BioLabs) and 5mL of Clearing Premix (PN 20300114).

### 3.3 Probe hybridization and MERSCOPE imaging

Tissue was then washed with Sample Prep Wash Buffer (3× at RT for 5 minutes), Formamide Buffer (1× at 37°C for 30 minutes) before hybridizing with our custom 721-gene MERSCOPE gene panel (100µL) and incubated at 37°C for 36–48 hours after clearing. After probe hybridization finished, samples were washed with Formamide Buffer (2× at 47°C for 30 minutes), Sample Prep Wash Buffer (2× at RT for 5 minutes), and stained for DAPI and PolyT (at RT for 15 minutes on a rocker). The samples were washed with Formamide Buffer (at RT for 15 minutes on a rocker), Sample Prep Wash Buffer (1× at RT for 5 minutes), and loaded into the MERSCOPE for imaging.

### 3.4 Data preprocessing

Raw MERFISH spots were decoded using the integrated MERSCOPE software (v233). Cell segmentation on the MERFISH output data was done using the Cellpose 2.0 algorithm [60] on the DAPI and PolyT signal of the median plane of imaging (4th out of 7), utilizing the Vizgen Post-processing Tool (https://github.com/Vizgen/vizgen-postprocessing). The segmented cell boundaries of the median imaging plane were then propagated to all other layers. A set of filtering parameters were applied to the detected cells to remove false positives and badly segmented cells. First, all cells with volume inside the population’s first and last 2% quantile were removed. For each cell that passed the volume filter, the maximum count of the blank probes (98 total blanks) was selected as the false detection threshold, and genes that have their detected count under this threshold were removed from the cell. Cell transcripts were then normalized to the maximum cell volume of the population, assuming a linear scale, before removing cells with the normalized total transcript count in the first or last 3% quantile or with less than 30 unique genes (excluding blanks). Lastly, cells that passed the filters had their transcripts data reset to the original non-normalized counts for downstream analysis.

Tissues were manually aligned to the Common Coordinates Framework v3 (CCFv3) [61] by matching the DAPI and PolyT histology stains to the reference atlas (25-µm resolution) in QuickNII [62]. Non-linear refinements to the registration were done using VisuAlign (https://www.nitrc.org/projects/visualign) to get the final indexed mask of CCFv3 labels for the registered tissue. The label mask was then fitted to filtered cells’ coordinates to transfer the CCFv3 labels to corresponding cell identities. Only cells labeled as Primary Somatosensory (S1, regardless of subregion) were used in downstream analysis (figure 3).

**Figure 3.**
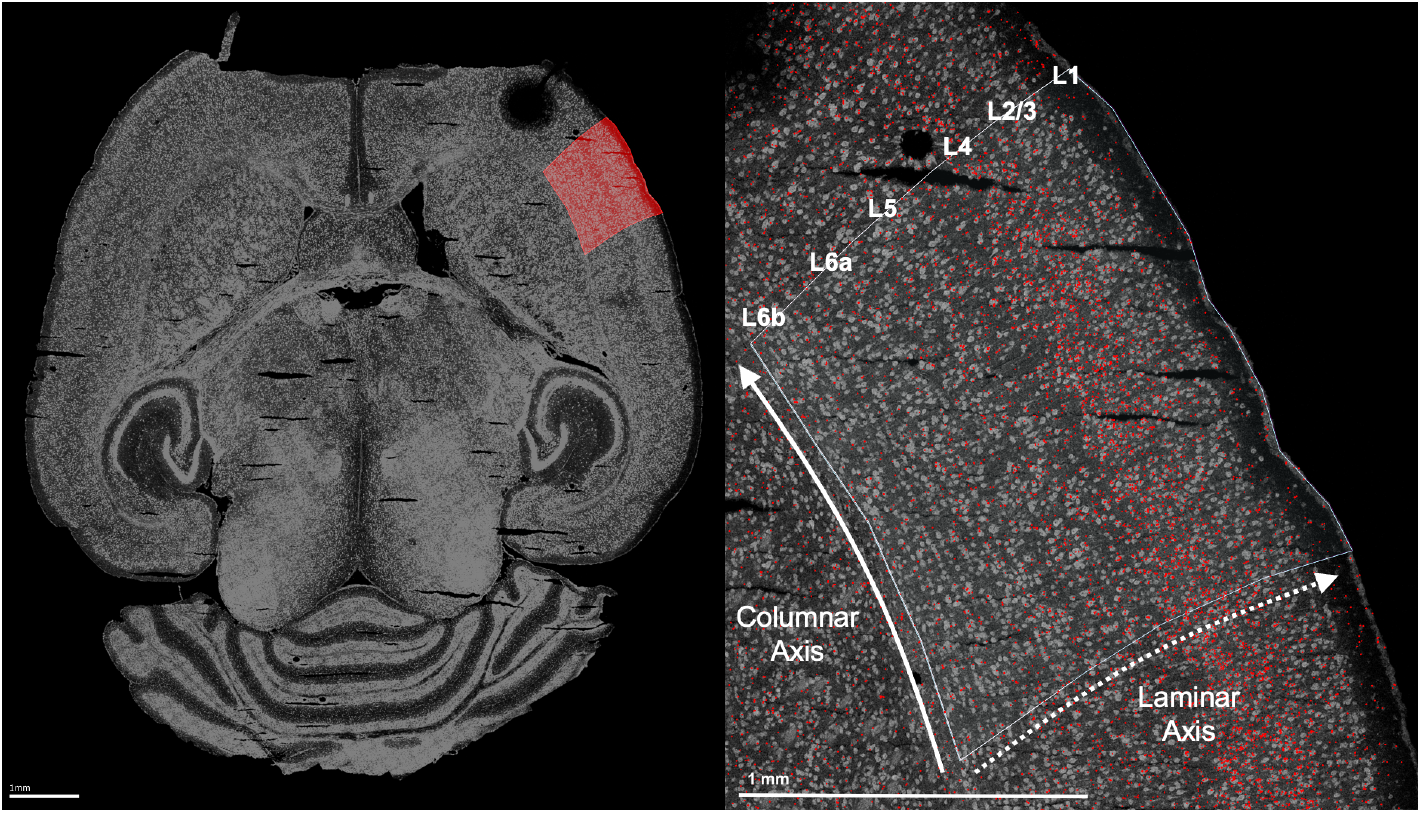
Horizontal slice from a P18 male wild-type mouse used in the analysis (mouse 4). Both visualizations produced via rendering of the transcripts observed on the MERFISH run. Left: Entire slice, right primary somatosensory cortex highlighted in red. The hole above highlighted area is a registration mark. Right: The same ROI, annotated with arrows showing the laminar and columnar axes with layers labeled. Red dots are RORβ transcripts observed on the MERFISH run. Note the higher transcript density in L4.

### 3.5 Transformation into laminar coordinates

After registration, we had a list of cells annotated with (1) coordinates ⟨**x, y**⟩ from cellpose giving the position of the cell’s center, (2) layer identity and ROI from the CCFv3, and (3) for each gene in our panel, a raw (i.e., non-normalized) count of transcripts in that cell. (So, each transcript was not assigned its own actual observed coordinate, but the coordinate of its containing cell.) The final preprocessing step was to transform the cellpose coordinates into a coordinate system defined by functionally meaningful axes. As we were interested in the laminar axis, we performed a series of geometric transforms which resulted in one axis perpendicular to the laminar boundaries identified by the CCFv3 registration and a second orthogonal axis presumably perpendicular to the cortical columns of S1 (figures 3, 4). These transformations were performed individually on each hemisphere of each mouse. Note that we excluded L1, as it contains very few cell bodies (figure 3). Each transformation was designed (and manually checked) to respect orientations, so that the origin was the deepest point perpendicular to the cortical surface (i.e., the deepest point in L6b) and also the most posterior point. Hence, the columnar axis ran from posterior to anterior and the laminar axis ran from L6b to L2/3.

**Figure 4.**
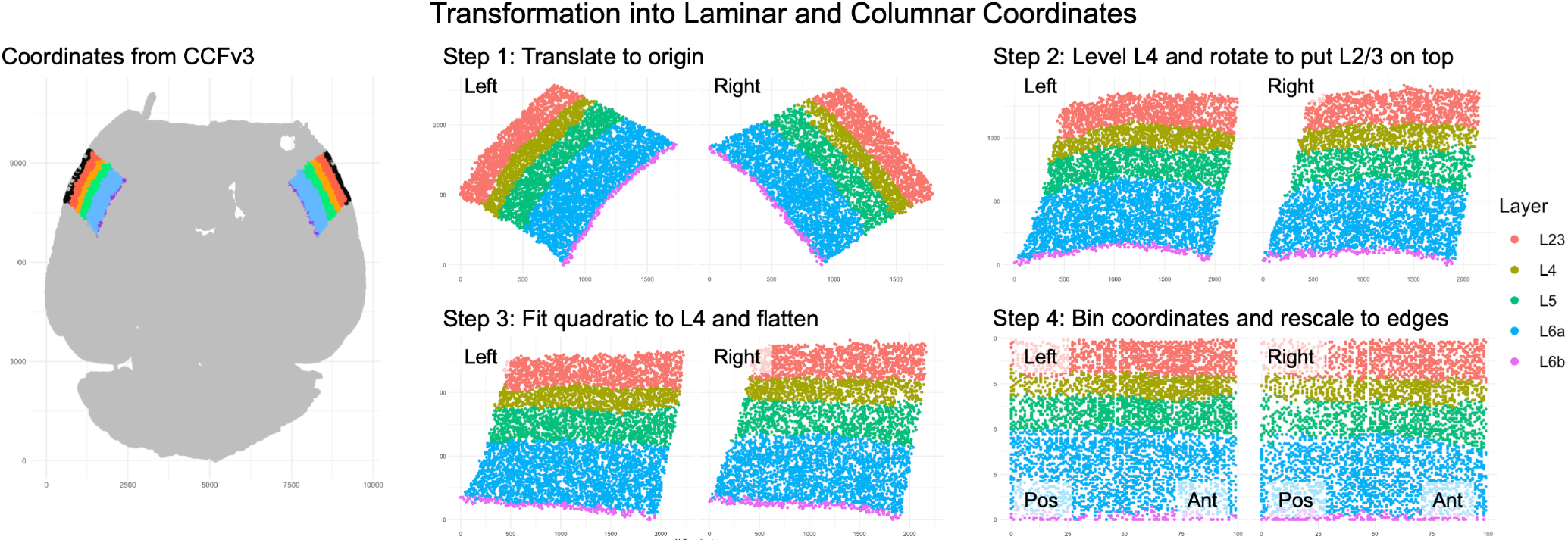
Transformation from post-registration cellpose coordinates into binned laminar and columnar coordinates. Data from mouse 4 used here as an example. Note that the displayed coordinate axes may not represent the true origin for the step, as each step often involved several trivial linear transformations to put the coordinates into a more convenient form. All steps shown with the left and right tissue patches equivalently positioned in the positive coordinate quadrant.

As shown in figure 4, there were four major steps to the coordinate transformation. (Step 1) The tissue patch was translated to the origin by subtracting the mean point of L5 along both axes. (Step 2) The tissue patch was leveled by fitting a linear model **y** = **mx** + **b** to the points of L4, then applying a matrix transform performing a rotation of arctan(**m**) radians (or its negative for negative slope). Coordinates were reflected across the horizontal axis, if necessary, to ensure L2/3 was on top. (Step 3) The curvature of the layers was flattened by fitting a quadratic model **y** = −**c**(**x** −**a**)^2^ + **b** to L4 using nonlinear least squares, then each point was transformed via ⟨**x, y**⟩ → ⟨**x, c**(**x** −**a**)^2^ + **b**⟩. (Step 4) The coordinates of both axes were binned so that the deepest, posterior most point was in bin ⟨0, 0⟩ and the most superficial, anterior point was in ⟨*b*_*p*_, *b*_*p*_⟩, then coordinates were “stretched” to fill the bins by, for each bin **x**, finding the max bin **y**_max_ holding a cell and rescaling the **y** component of all points in **x** with **y** ≥ *b*_*p*_*/*2 so that *b*_*p*_*/*2 → *b*_*p*_*/*2 and **y**_max_ → *b*_*p*_ − |*ϵ*|_abs_, for some small jitter *ϵ*. This rescaling was weighted by each cell’s distance from the midpoint *b*_*p*_*/*2, so that points near the midpoint only received a small fraction of the rescaling while those at the max receive all of the rescaling. An analogous rescaling was performed for points in **x** such that **y** *< b*_*p*_*/*2 to pull **y**_min_ down to zero (plus some positive jitter) and two analogous rescalings were done for the **y** bins so that **x**_max_ was pulled over to *b*_*p*_ and **x**_min_ was pulled back to zero. The exact algorithm for these steps can be found in the wispack code. After performing this transformation, the final binned coordinates *x* used to fit the WSP model were obtained by simply dropping the **x** coordinate (i.e., the coordinate orthogonal to the laminar axis) so that *x* = **y**.

## 4 Results

### 4.1 Model fit

After preprocessing the data from four mice, we had 54,116 cells in S1 containing 341,184 transcripts from a total of six genes *g* (table 1). While there was substantial variation across mice (table 3), it was relatively balanced across ages. As explained above, no normalization was performed on these counts.

**Table 3:** Age, cell counts, and mean transcript count per gene per cell for each mouse. The asterisk ∗gives the mean across the entire 721-gene panel, while the far right column gives the mean across only the six genes modeled here. All counts for the analyzed ROI (S1) only.

| Mouse | Age | Cells | $\mu(y/g^*/\text{cell})$ | $\mu(y/g/\text{cell})$ |
| --- | --- | --- | --- | --- |
| 1 | P12 | 13,237 | 1.677 | 1.290 |
| 2 | P18 | 16,954 | 1.884 | 1.238 |
| 3 | P12 | 10,444 | 1.029 | 0.650 |
| 4 | P18 | 13,481 | 1.212 | 0.891 |

Coordinates along the laminar axis of each tissue sample were divided into *b*_*p*_ = 100 bins. Mouse number was used as the levels *k* of the random-effect factor *ν*. There were five random-effect levels all together, the four mice plus the pseudo reference level. Hemisphere and age were the two fixed-effect factors analyzed, i.e., *ξ* = ⟨*ξ*_1_ = hemisphere, *ξ*_2_ = age⟩, with the left hemisphere and P12 being used as reference levels and the right hemisphere and P18 treatment levels. Thus, there were four treatment interaction levels *w*: (1) baseline (left, P12), (2) hemisphere (right, P12), (3) age (left, P18), and (4) the age-hemisphere interaction (right, P18). Six grouping variables (genes, *g*), five random-effect levels (mice plus extrapolated reference, *ν*), four treatment interaction levels (hemisphere × age, *w*), and 100 position bins (*x*) gives 12,000 potential observations (rows of data) to fit. The count *y* for each row was gotten by summing the transcript count of that row’s gene *g* from all cells annotated by the values *x, ν*, and *w* of that row. (A similar summing of counts is performed by ELLA [63]; summing does not affect model fit because the sum of two Poisson-distributed variables is itself Poisson-distributed.) As each mouse could only be observed at one age, 4,800 of these rows were empty (half of all rows not including the pseudo random-effect reference level), although the WSP model still made predictions for these empty rows.

The initial degree and parameter estimation by LRO change-point detection (equations 20–26) yielded 300 parameters to be fit with a boundary set of size 1,260 (equation 36). Next, dispersion factors *ζ* were estimated (equation 7) and the parameters were fit to the data using L-BFGS by penalized likelihood maximization (equations 30–39). The residuals from this fit are shown in figure 5 as a histogram and Q-Q plot. The posterior distribution P(*β, ρ*|*y, ζ*) of the WSP model parameters was estimated with both MCMC and bootstrapping, although only the samples collected with bootstrapping were used to compute confidence intervals and *p*-values. In both cases 10,000 samples were collected, via (1) a Metropolis–Hastings random walk for MCMC (equations 40–43), and (2) for bootstrapping by, for each data row (*x, ν*, and *w* combination) and gene *g*, randomly sampling (with replacement) cells from that row and summing the transcript counts for that gene, then refitting the model with L-BFGS to the resampled sums using recomputed dispersion factors *ζ*. Both sampling procedures were run on an Apple M2 Ultra with 128Gb of ram, single-threaded for the random walk and forked across ten threads for the bootstraps. Each full bootstrap resample and refit with L-BFGS used on the order of 10Gb of ram, presumably mostly due to Stan’s use of expression graphs to compute gradients.

**Figure 5.**
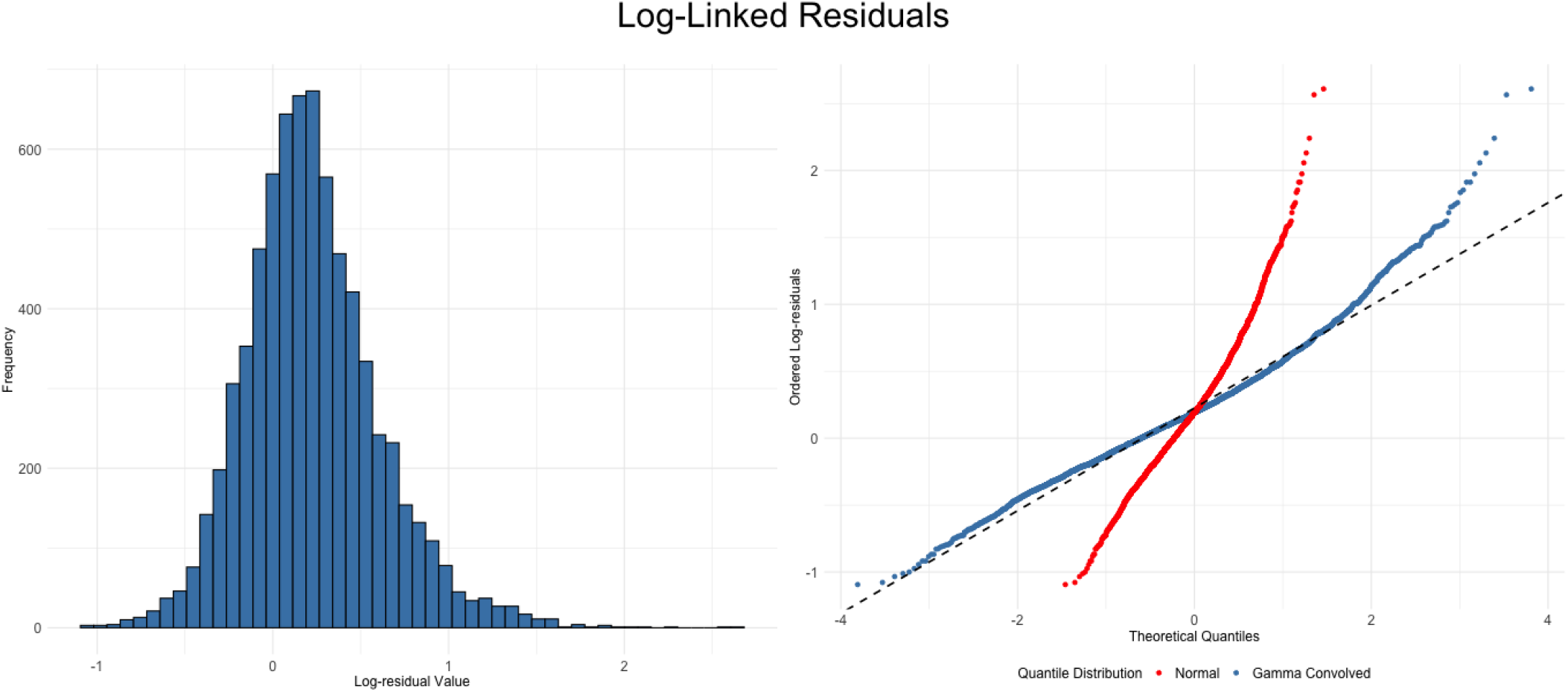
Histogram (left) and Q-Q plot (right) for the residuals from the L-BFGS WSP model fit. The Q-Q plot shows both the theoretical quantiles from a normal distribution (red) and the theoretical quantiles derived from a normal distribution convolved with the gamma distribution used to fit the data (blue, equation 31). The way the blue points are closer to the dashed black line than the red points shows that the residuals are more normal when the over-dispersion from the gamma distribution is taken into consideration. That is, modeling the over-dispersion with a gamma kernel notably improves model fit.

Total time to complete the random walk was 42 minutes. We left a gap of one accepted step between saved accepted steps in order to “thin” the sampling and minimize next-step autocorrelation, so the walk effectively took 20,000 steps (averaging 0.124 seconds per accepted step). However, despite trying many step sizes, priors, and gap sizes, observed autocorrelation was extremely high (figure 6F). In addition, the MCMC samples were not as normal as the bootstrap samples, as indicated both by lower Shaprio-Wilk *p*-values (figure 6E) and more parameters with multimodal distributions (figure 6D). The nonideal distributions and high autocorrelation of the parameter samples from the MCMC walks were due to a failure to large drifts and a failure to stabilize around a mean (figure 6A, B). The MCMC algorithm did find a stable negative log-likelihood value, however that value was still notably higher than the one found by L-BFGS (figure 6C). We interpret these results as suggesting that the likelihood landscape is rocky and that the MCMC random walk is hopping between local minima in parameter subspaces. While the acceptance rate on the run reported here was low, at 0.133, the many walks observed in development with more ideal rates of 0.2–0.3 still exhibited these same issues.

**Figure 6.**
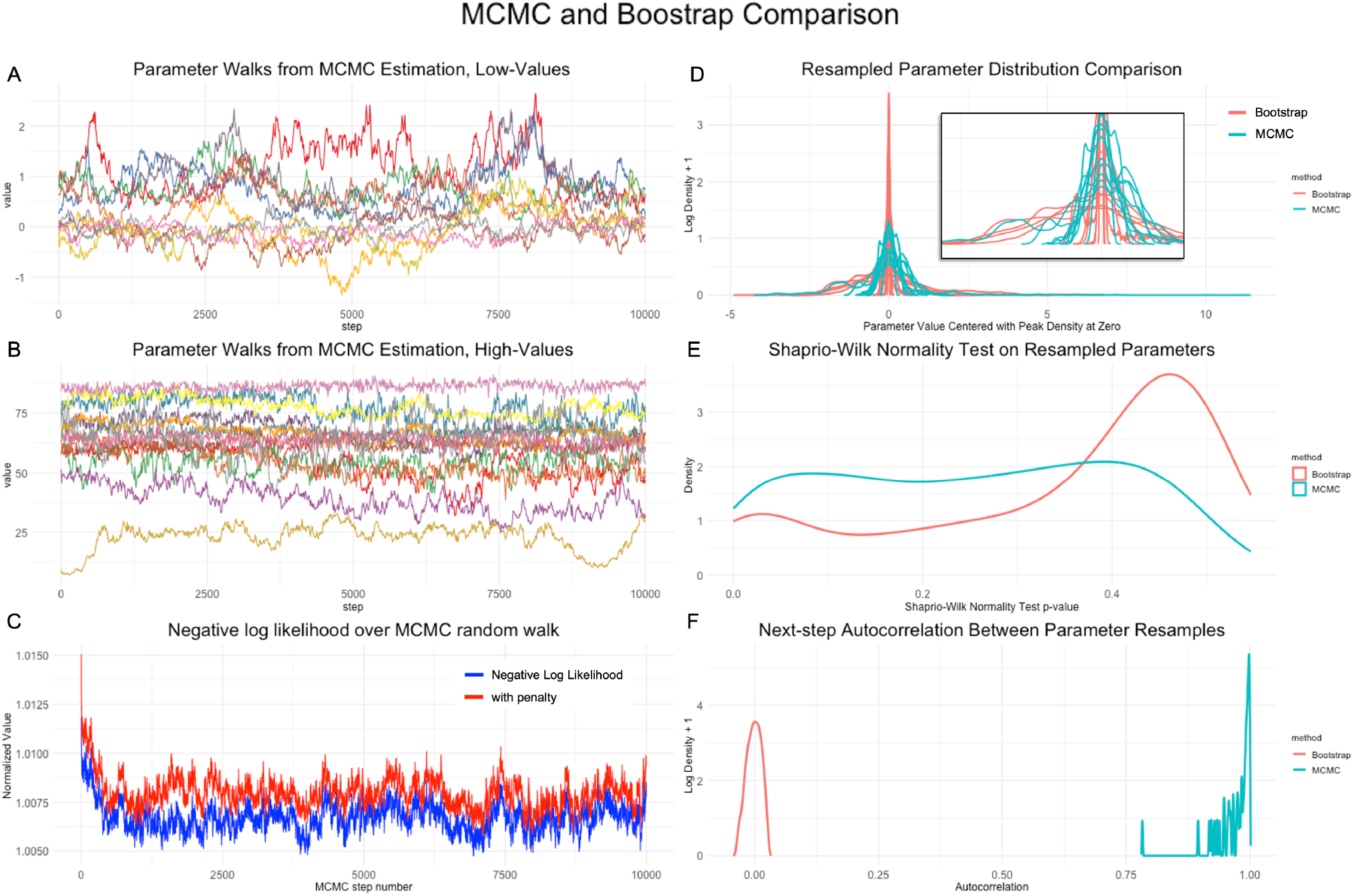
Diagnostic plots for MCMC parameter samples. (A) Ten randomly selected low-value (*<* 20) model parameters, plotted over the 10,000 step random walk. (B) Same, but for all high-value (≥ 20) model parameters. (C) Negative log-likelihood and penalized negative log-likelihood over the walk, normalized relative to the L-BFGS fit value. (D) Distribution of ten representative model parameter samples, for each of the MCMC and bootstrap methods. (E) Density of the estimated Shaprio-Wilk *p*-values across all model parameters, as sampled by both MCMC and bootstrapping. (F) Distribution of the correlation between parameter sample *I* and sample *I* + 1, as sampled by both MCMC and bootstrapping.

**Figure 7.**
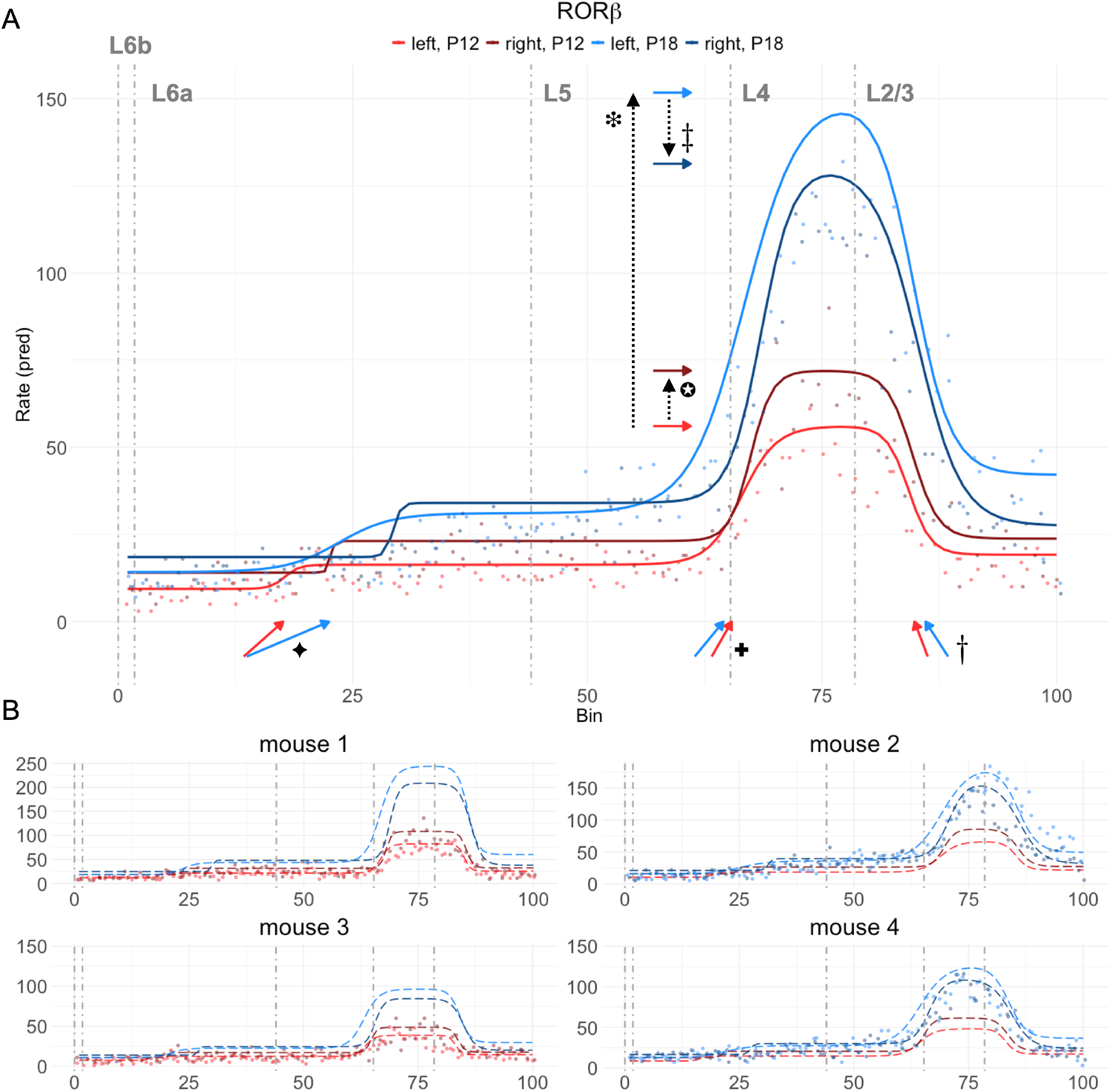
WSP model fitted to RORβ transcript counts across the laminar axis. Dot-dashed vertical grey lines show the laminar boundaries estimated independently by CCFv3 registration. Points are transcript counts in each *x* bin. (A) Interpolated data (points) and predictions (solid lines) for RORβ with no random effects. Colored arrows along the *x* axis point to the locations of the three transition points at age P12 (red) and P18 (blue), with the angle of their tails showing the transition point gradient. Colored horizontal arrows just left of L4 point to the predicted rate value of block 3 in each of the four conditions. Dotted black arrows show direction of associated effects. Ornamental glyphs by each arrow refer to lines in table 4 giving the estimate for the associated effect. (B) Data and predictions for RORβ for the individual mice. Points show data collected from the mouse, dotted lines show both the model predictions for that data and the interpolated model predictions for the unobserved age.

Total time to complete the bootstraps was 84 minutes, averaging about 0.51 seconds per bootstrap, implying about 5.1 seconds per thread. L-BFGS converged (tolerance of 10^−6^) on all bootstraps (median of 182 iterations, max of 266), giving 10,000 samples of the parameter vector Φ, plus the original parameters fit to the full data. These 10,001 parameter samples were used to compute 95% confidence intervals on all 300 parameters (equation 44) and *p*-values for fixed-effect parameters *β* not including the baselines (171 parameters total, equation 47).

### 4.2 FSE tests

With one exception, the estimates of transition points between expression rates made by the WSP model were near the laminar boundaries obtained independently from the CCFv3 registration, as can be confirmed visually by comparing the plots in figure 8 with the expected layers listed in table 1. The exception is SATB2, which appears equally distributed across all but the deep part of L6, while we expected significant upregulation in L2/3. However, this distribution is clearly visible in the data itself (figure 8), and therefore does not reflect any obvious failure of the model to the fit the data. In addition, sharp upregulation of SATB2 in L2/3 is most well established in embryonic development [32] with less known about the later ages (P12, P18) for which we have data.

**Figure 8.**
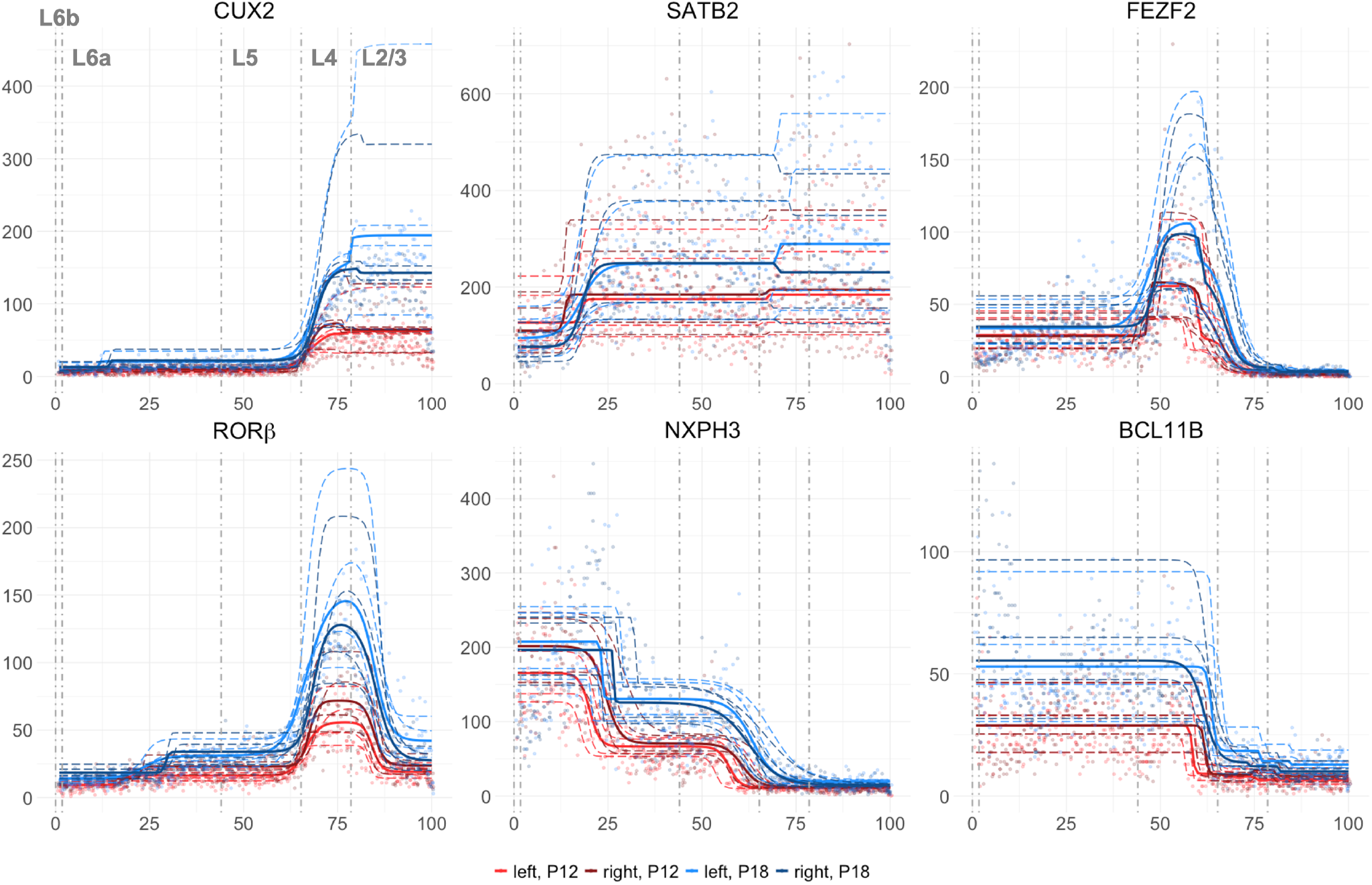
WSP model fitted to transcript counts across the laminar axis of S1, all modeled genes. Data and predictions plotted just as in figure 7, except that extrapolated values without random effects and values for individual mice are combined on a single plot. Points are data, solid lines show predictions with no random effects, dashed lines show predictions for each random effect level (i.e., each mouse).

Consistent with our hypothesis, we found effects of both age and laterality on RORβ expression, including two potential FSEs of note (figure 7). (1) The WSP model showed an increase in RORβ expression from P12 to P18 (the two ages for which we have data) across all but the deep part of L6 along the laminar axis, with the largest natural-log fold change in block 3, the block most closely aligned with L4 (*β* = 0.985, *p <* 0.001). (2) The WSP model showed a significant effect of laterality, both at the reference age of P12 and as an interaction with the treatment age of P18. Interestingly, these effects went in opposite directions: RORβ expression in block 3 in the right hemisphere was increased (relative to the left) at P12 (*β* = 0.246, *p <* 0.001), but decreased (relative to the left) at P18 (*β* = −0.389, *p <* 0.001). Although potential FSEs on RORβ expression were limited to effects on rate, numerous significant effects on slope scale and transition point location were observed in the other modeled genes (table 4).

**Table 4:**
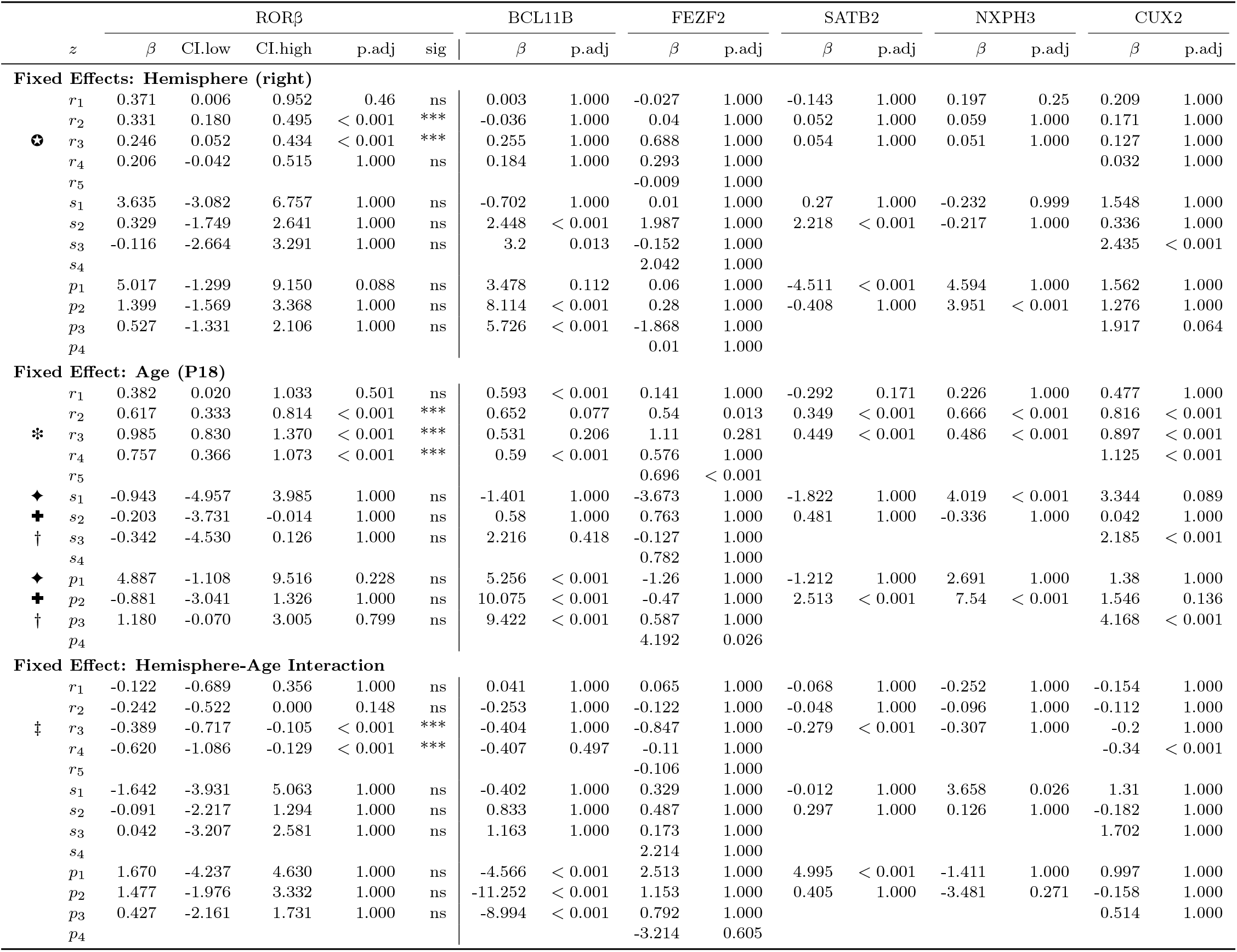
Fixed effect estimates *β* given as natural-log fold changes, *p*-values and 95% confidence intervals adjusted with Holm-Bonferroni correction. Ornamental glyphs refer to arrows in figure 7 which visualize the effect.

## 5 Discussion

Despite a growing number of new tools for analyzing ST data, before WSP models there were none available for answering seemingly simple questions such as whether RORβ expression rate is lower in the right hemisphere compared to the left. While technically possible, tackling such a question would have required bespoke data preprocessing with a high risk of biasing the answer, e.g., by manually specifying the intracortical boundaries of RORβ expression. WSP models simplify this preprocessing to a basic step — aligning spatial coordinates with an axis of interest — with less risk of biasing the answer. WSP models enable this less-biased hypothesis testing by offloading the estimation of key spatial parameters (especially transition points between expression rates) to a data-driven, likelihood-based regression. Thus, WSP models provide a way to test hypotheses about FSEs, i.e., about the factors which have functionally relevant effects on the spatial distribution of gene expression. By enabling testing of FSEs, WSP models provide a powerful new tool for uncovering the genetic mechanisms behind biological and cognitive functioning.

### 5.1 FSEs in the whisker barrel system

To demonstrate WSP models’ capability to detect functional spatial effects, we hypothesized that RORβ expression in developing mice varies significantly with age and hemisphere. We hypothesized an effect from age based on prior work showing two key developmental trends in mice [10, 11]: (i) barrel distinctness and RORβ expression rise in parallel from birth P0 to P7, and (ii) barrel distinctness then wanes from P10– P30, with a significant decrease by P20 in males. These observations suggest a tight early correlation between RORβ and the physical refinement of barrels, yet leave open whether that relationship persists later. Results from the WSP model, fit to data we collected from P12 and P18 male mice, provides a preliminary answer (table 4, figure 7). Interestingly, instead of the predicted decline, the WSP model showed that RORβ increased further at P18 (compared to P12), even as barrel distinctness presumably fell, revealing a decoupling of gene expression from the macroscopic barrel phenotype in late development. This late rise implies that RORβ’s role extends beyond initial barrel sharpening, underscoring WSP models’ power to pinpoint age-dependent shifts in gene-circuit relationships.

Laterality is essential for a variety of sensorimotor tasks [64], but when and how it emerges in cortex is still unclear. We recently showed circuit-level mechanisms that could drive left–right specialization in the mouse auditory cortex [30], yet molecular programs remain unresolved. Observations of laterality during development have also been observed in S1 of rats. At P20, total cortical area containing barrels was observed to be significantly greater on the left (and therefore smaller on the right) in males [31]. The WSP model fit to our data from P12 and P18 male mice showed comparable molecular asymmetry (table 4, figure 7). At P18, RORβ expression was significantly lower in the right hemisphere than in the left — a pattern that matches the smaller right-hemisphere barrel field reported in P20 male rats. However, the WSP model also showed a strong interaction with age. RORβ expression increased from P12 to P18 in both hemispheres, but increased significantly more in the left. Indeed; the WSP model showed that RORβ expression at P12 was higher in the right, opposite the pattern observed in P20 male rates. Although some adult mouse studies (*>*P60) report no barrel-field laterality [65, 66], those analyses may miss the transient window we and others observe in pups (*<*P20).

### 5.2 Comparison with SVG testing

Recent statistical methods for ST data have focused on detecting spatially variable genes (SVGs), both within a single cell and across larger tissue samples [25, 26]. SVG detection methods include both artificial neural network classifiers and traditional regression models. Most regression approaches fit a model to transcript counts, although some fit to a proxy of spatial gene count distribution, such as diffusion time (sepal [67]) or interaction energy (BOOST-MI [68]). Many of the count-based regression approaches (e.g., SpatialDE [69], SPARK [70], SpatialDE2 [71], SMASH [72]) predict gene expression rate (a log-linked raw transcript count or normalized transformation of it) with a generalized linear model, under the assumption that observed counts are Poisson distributed, possibly also using an inner convolution to handle over-dispersion (equations 4, 5). These approaches further assume that one of the terms in the model, either some set of covariates *X* or coefficients *β*, comes from a multivariate normal distribution (MVN). The idea is that if the covariance matrix, or *kernel*, of this distribution is a function of spatial position, then this spatial covariation indicates that the gene being tested is spatially variable.

The main difference between these spatial kernel-based SVG tests and WSP models is the latter’s emphasis on theory-driven hypothesis testing. While some SVG tests are explicitly formulated as tools for hypothesis testing, they face one or both of two limitations [26, 73]. The first limitation is that these tools only allow for testing the hypothesis that a gene is spatially variable. For example, SPARK [70] models expression rate Λ as a linear function of position *x* including a term **b**(*x*) ~ MVN(0, *τ* **K**(*x*)) assumed to come from a multivariate normal distribution centered on zero with covariance **K**(*x*) scaled by *τ*. (The symbols “*τ* “, “**b**”, and “**K**” are their notation.) The Satterthwaite method with score statistics is used to compute a *p*-value for the null hypothesis that *τ* = 0. While this procedure tests for non-zero spatial covariance (i.e., tests for SVGs), it does not allow for testing for effects on that spatial covariance.

SPARK also exemplifies the second limitation common to SVG tests (e.g., SpatialDE, MERINGUE, SpaGCN, DESpace): it does not allow for testing for *between-group* effects [73]. For example, SPARK cannot be used to test whether a factor like cell-type, age, or gene-knockout affects the spatial-covariance scalar *τ*, i.e., affects whether a gene is a SVG. SPARK is limited to *within-group* testing, i.e., to testing whether a gene as observed in one set of conditions (e.g., holding cell-type and age constant) is spatially variable. While between-group testing has been a staple of traditional differential expression tools such as DESeq2 and some recent SVG tools such as SPADE [73] have introduced it as well, the first limitation remains.

These limitations are largely intrinsic to the structure of common SVG methods, which essentially work by testing for non-zero spatial covariance (figure 1A). The focus on spatial covariance is a serious hurdle for integrating SVG tools with genomics databases such as GO [23] and KEGG [24], as even with the addition of between-group testing, effects on spatial covariance are not readily interpretable as log fold changes. Further, it is unclear how to extend these methods into tests for FSEs. As articulated in the difference between equations 2 and 3, testing for FSEs requires an explicit parameterization of the spatial distribution. A FSE is an effect on some parameter of the spatial distribution of a gene. Although one recent SVG tool (ELLA, [63]) moves away from modeling spatial covariance to explicit modeling of spatial distribution, its mathematical representation of spatial distribution is not in terms of easily interpretable parameters, such as transition-point location and slope. This would make it challenging to use it to test for FSEs.

### 5.3 Random variance estimation

Another difference between WSP models and SVG tests is that the latter offer no way to estimate random effects from variance due to differences between individual tissue samples. This issue is especially pressing, given that most ST studies use small tissue-sample sizes (*n <* 5), including our demonstration here. The need to estimate random effects in transcriptomics data (both spatial and RNA-Seq) is most acute as it relates to testing for effects from age and other temporal factors such as rearing conditions.

The problem is that temporal factors are of theoretical interest and can be expected to have systematic (i.e., fixed) effects, but current technology usually does not allow for sampling multiple time points from the same individual animal. Thus, researchers attempting to model transcriptomics data from multiple time points will need to account for the effects of time, but usually have no direct way of experimentally separating temporal effects from random variation between samples. A sufficiently large sample at each time point will allow for estimating whether there is age-related variance that is unlikely given the individual variation within an age, thus solving the problem. However, ST data collection in particular is still extremely costly in terms of both money and time. Thus, more indirect strategies will be necessary for the foreseeable future.

Mixed-effects modeling, i.e., modeling individual variance explicitly through random-effects terms *ρ*, is one such indirect strategy [45]. Along with other strategies (e.g., regularization or “shrinkage”), mixed-effects modeling has been used to account for random effects by popular tools such as DESeq2 [21]. However, mixed-effects modeling is helpful only if the model is sufficiently constrained. Holding random effects constant across grouping variable levels (i.e., genes) is one way to obtain the needed constraint. However, we felt it likely that random variation between individuals would depend on the specific gene, and so gave each random level a random effect value for each gene. With WSP models, the needed constraint comes instead from including a second factor (e.g., hemisphere) which can be sampled fully within each random level and is orthogonal to, and interacting with, the problematic factor (e.g., age) which can only be sampled once per random level.

A second way to address the problem is with priors. Instead of fitting the model by maximizing the likelihood of the observations given the parameters, the model could be fit by maximizing the product of the likelihood and a prior for the given parameters, which would be equivalent to maximizing the posterior for the parameters. Potential priors could take many forms. For example, the assumption of a normal effect distribution could be codified into a prior by setting

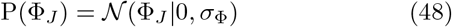

Another approach would be to estimate an expected value *R* for the ratio of total variance over random variance from the observed counts and set

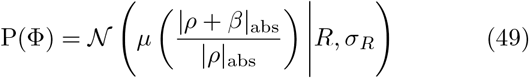

A third approach would be to estimate expected random variation *V* based on prior data and set

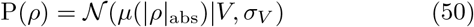

Future versions of WSP models could incorporate one or more of these priors to improve random variance estimation. More broadly, we hope that WSP models raise awareness of the need to handle the small sample sizes in ST data.

### 5.4 Limitations and future development

WSP models have limitations worth addressing in future development. First, parameter estimation takes substantial time and compute power, even for a few dozen genes and a handful of tissue samples. While wispack is already highly optimized, fitting WSP models in 5–10 seconds per thread, compared to fits during early develop with the R optim function which took over twenty minutes, we estimate that improved optimization methods and more efficient code could get fit times down into the 1–3 second range. However, we don’t believe fit time is a critical issue, as WSP models are meant for theory-driven hypothesis testing. Large-scale data exploration is not feasible, but also is not the intended use.

A second limitation is the need for bootstrapping. While MCMC is possible, we have so far seen mostly poor performance (figure 6). We assume this is due to the many exponential nonlinearities involved in a WSP model and the complex parameter boundaries. These aspects of WSP models presumably lead to a rocky likelihood landscape. A random walk may not be suitable for navigating this landscape; a directed method such as gradient descent may be needed. The downside of bootstrapping is not only the increased computation time, but also the need for data that is structured in a way allowing for resampling. Improvements to step-size selection (equation 41) or an alternative to Metropolis–Hastings sampling may make MCMC parameter estimation more reliable. The constraint to a single spatial dimension *x* is also a limitation. First, there may be functionally relevant expression distributions over two or more dimensions. Second, the need to align to a single spatial axis of interest both necessitates bespoke coordinate transformations (e.g., figure 4) and leaves room for biased axis selection. Future work on WSP models should aim to generalize the mathematical principles to two or more dimensions. A related limitation is the need to handle time (e.g., age) as a two-level factor. We hope to incorporate an explicit time-like dimension into future iterations of WSP models.

Statistically, the assumption of gene independence in the likelihood calculation (equation 30) is a limitation and the simplistic estimation of dispersion factors (equation 7) could be improved. The former is a major limitation, given that often one will want to test entire pathways of causally related genes. We see the latter as a minor limitation related to refining the best possible fit. While improvements in fit are always desirable, we believe there is more to be gained from improving the LRO change-point detection algorithm for degree estimation over improving the dispersion factor estimates.

A final question concerns the extent to which the rate, slope scalar, and transition point location parameterization is itself a limitation. At least in the case of a single dimension, we think it’s unlikely that there are any biologically realistic transcript distributions which cannot be described by the parameters of a WSP model. However, even so, it’s an open question whether there are hypotheses this parameterization cannot test, e.g., because those hypotheses concern incommensurate distribution parameters. This question can be settled only after seeing the hypotheses researcher attempt to test with WSP model.

## 6 Conclusion

We developed a novel form of logistic regression based around a parameterization for spatial distributions which allows for both random-effect estimation and between-group hypothesis testing with spatial transcriptomic data. To demonstrate the potential of WSP models, we tested for effects of age and laterality on the distribution of RORβ expression across the laminar axis of S1 in mouse pups, which is known to play a critical role in the development of cortical whisker-input organization. The results (1) suggest that the mechanisms behind this development are age and hemisphere-dependent and (2) underscore the utility of WSP models for uncovering subtle molecular asymmetries and testing hypotheses related to biological and cognitive function.

## Appendix

### A.1 Wispack MWE

To install wispack, clone the repository at https://-github.com/michaelbarkasi/wispack.git and follow the install instructions in the readme file. The minimum input required to use wispack is a data frame in the format used by standard R functions for linear modeling. For example, when modeling data with the formula `observation _ factor1 * factor2`, standard functions like lm expect a data frame with columns named `observation, factor1`, and `factor2`, with each row being an observation. Calling the wisp function with such a data frame will fit a WSP model to the data, estimate confidence intervals and *p*-values with a MCMC walk, and output a number of data frames and summary plots with results. For example, the data (post transformation into laminar coordinates) used in this demonstration is provided in the git repository for wispack. The following code fits a WSP model to it with default settings:

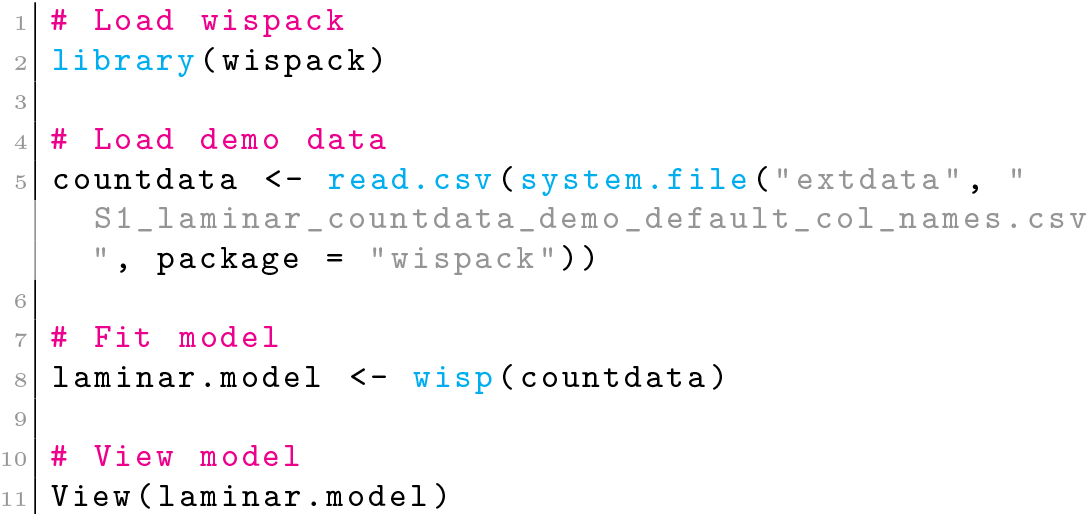

Unlike standard functions for linear models in R, `wisp` does not take a formula specifying a model, as the model formula is always of the same form (equation 10). What varies are the variables to use as fixed effects, random effects, etc. These are specified by giving the `variables` argument a list of elements naming columns in the data, each labeled with the model component given by that element (`count` for observed counts, `bin` for the spatial coordinate, ran for the random effect variable, etc):

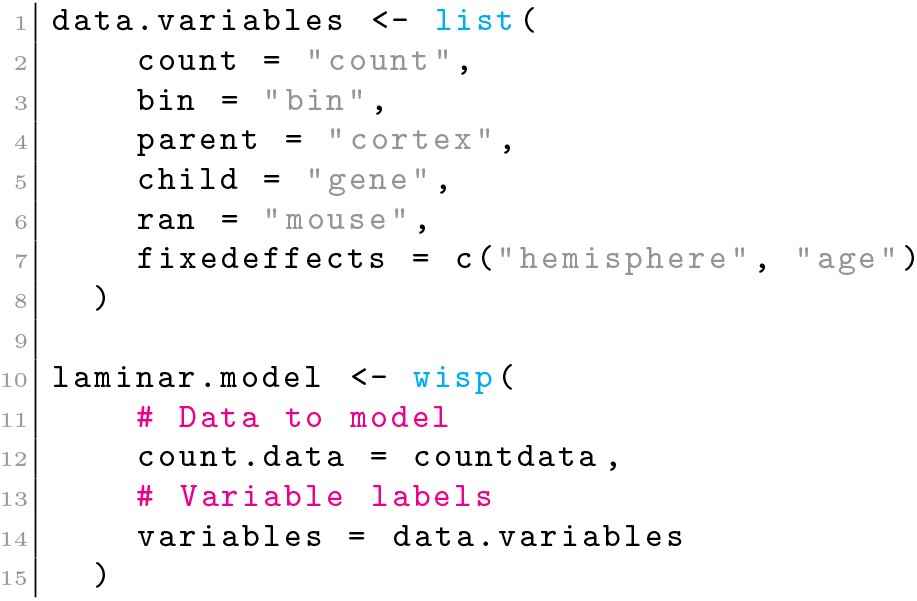

Note that although in this paper we only described a single level grouping variable (for genes), wispack is coded to handle two-tiered grouping (with a parent and child level), hence the `parent` and `child` arguments instead of a singular `grouping`. If no variables are specified, wispack looks for columns in the data with names matching the labels in the `data.variables` example list. There are many other settings, e.g., for controlling the MCMC walk, running bootstraps, adjusting the LRO algorithm, etc. These are all shown in the `full options.R` demo in wispack. The git repository accompanying this paper contains the code with the settings used in the demonstration reported here (https://github.com/Oviedo-Lab/wspmmmethods.git).

### A.2 Sigmoid rate theorem

**Theorem:** If *s*_*i*_ ≫ 0 for all *i* and *i*′ is the largest *i* such that *x* ≫ *p*_*i*_, then Ψ_*d*_(*x, r, s, p*) ≈ *r*_*i*_′_+1_.

**Proof:** By definition (equation 10):

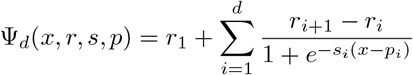

For all *i*, if *s*_*i*_ ≫ 0, the tendency of the exponential in the denominator of the summand to be either a very large value or a value near zero implies that:

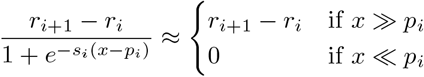

It follows that:

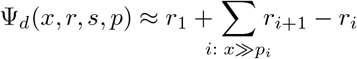

Using *i*′ for the largest *i* such that *x* ≫ *p*_*i*_, this can be rewritten as:

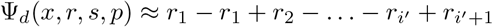

In the term on the right-hand side, all *r*_*i*_ cancel out except for the last, leaving Ψ_*d*_(*x, r, s, p*) ≈ *r*_*i*_′_+1_.

### A.3 Sigmoid slope theorem

**Theorem:**

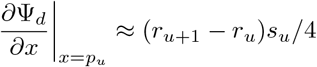

**Proof:** By the linearity of derivatives and definition of Ψ in equation 10:

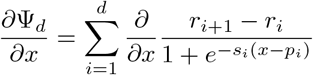

By the quotient rule and basic algebra, this equals:

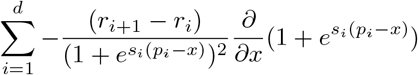

By the linearity of derivatives, the chain rule, and more algebra, this equals:

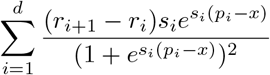

Evaluating this formula at *x* = *p*_*u*_ gives:

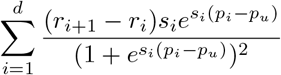

Note that for all *X, e*^*X*^ */*(1 + *e*^*X*^)^2^ ≈ 0 unless *X* ≈ 0. In our case, *X* = *s*_*i*_(*p*_*i*_ − *p*_*u*_), which only approaches zero if *s*_*i*_ ≈ 0 or *p*_*i*_ ≈ *p*_*u*_. The former is impossible, given the constraint on the slope parameter that *s >* 0. The latter is also impossible when *i* ≠ *u*, given the constraint that there exist a nontrivial buffer distance between all transition points *p*. Thus:

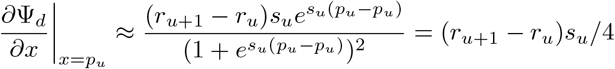

### A.4 Warping limit theorem

**Theorem:** Equation 18 is true:

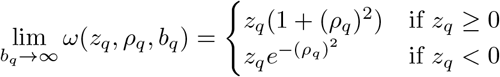

**Proof:** Begin by considering the limit of the warping ratio (equation 16) as the bound *b* approaches infinity:

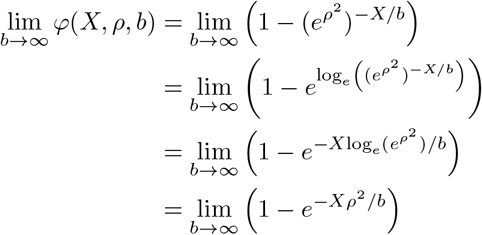

It is well known that:

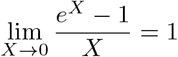

It follows that 1 − *e*^*X*^ ≈ −*X* as *X* → 0, and so:

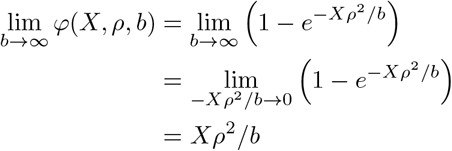

Now consider the full warping function for positive *ρ*:

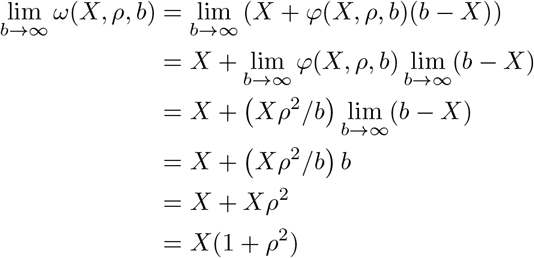

Next, consider the full warping function for negative *ρ*:

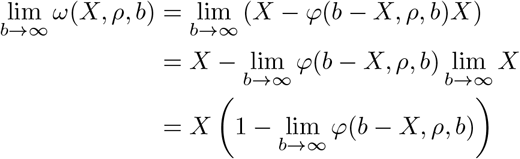

The original reasoning about the warping ratio’s behavior as the bound approaches infinity implies that:

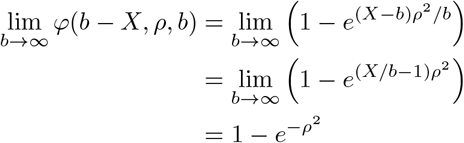

And so, finally:

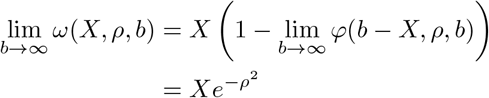

### A.5 Kernel convolution solution

Equation 31 gives the likelihood of the observed data, given the predicted kernel rate and estimated gamma dispersion. This likelihood is given by integration over a Poisson-gamma convolution:

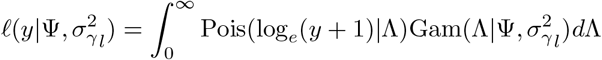

This integral can be solved analytically. Note that the above equation parameterizes the gamma density function in terms of its expected value and variance. In order to use the standard formula for gamma density, these need to be converted to the standard parameterization in terms of “rate” (rt) and “shape” (sp), by setting 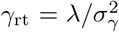 and 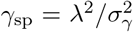. Using standard formulae for the Poisson and gamma density functions:

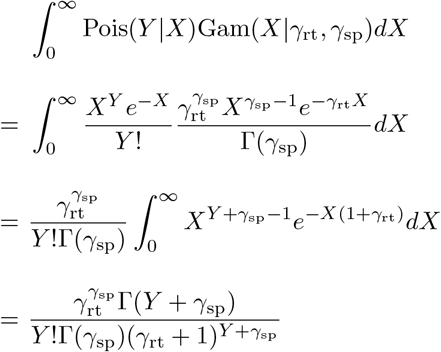

For generalization to the reals, wispack replaces *Y* ! with Γ(*Y* + 1). For numerical stability, the difference between the sum of the natural log of the numerator terms and the sum of the natural log of the denominator terms is computed first, before returning the exponential of this difference.

### A.6 The rate-boundary theorem

**Theorem:** Ψ *>* 0 if and only if *r*_*i*_ *>* 0, for all rate components *r*_*i*_ used in the calculation of Ψ.

**Proof:** The left-to-right direction of the biconditional follows immediately from the sigmoid rate theorem (appendix A.2). The right-to-left direction does not also immediately follow because the sigmoid rate theorem does not constrain what happens to Ψ near the transition points *p*. So, consider the case when *x* ≈ *p*_*i*_′. In this case:

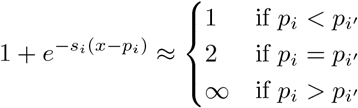

Thus, equation 10 can be rewritten as:

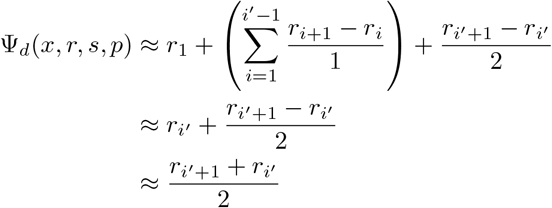

Assuming that all rate components *r*_*i*_ *>* 0, it follows immediately that Ψ_*d*_ *>* 0. Indeed, as expected, when near a transition point, Ψ_*d*_ is approximately the mean of rates of the blocks immediately before and after that point.

### A.7 Likelihood ratio computation

Equation 22 defines the log-likelihood ratio *L*_ratio_ for single elements 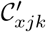 in the log-linked count matrix 𝒞′ as follows:

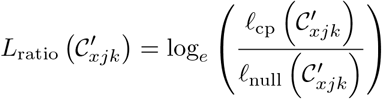

To compute this value, we use the quotient identity:

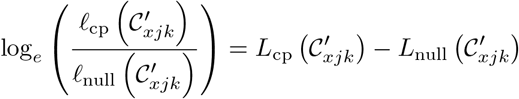

To compute *L*_cp_ and *L*_null_ for a point *x* in column *jk*, let 𝒳be a window of points centered on *x*. This set of points defines a set of log-linked counts 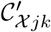:

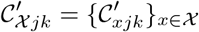

The log-likelihood of the null hypothesis that there is no change-point is defined:

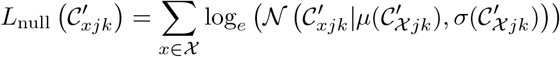

For each window 𝒳, the log-likelihood that the window contains a transition point at its center can be computed by summing the log-likelihoods of the null hypothesis for the points *x* in the first half 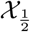 and second half 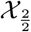of the window:

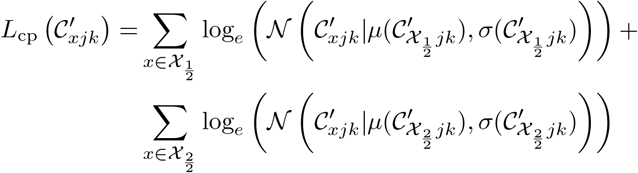

